# SUPT3H-less SAGA coactivator can assemble and function without significantly perturbing RNA polymerase II transcription in mammalian cells

**DOI:** 10.1101/2021.07.09.451791

**Authors:** Veronique Fischer, Elisabeth Scheer, Elisabeth Lata, Bastian Morlet, Damien Plassard, Stéphane D. Vincent, Dominique Helmlinger, Didier Devys, László Tora

## Abstract

Coactivator complexes regulate chromatin accessibility and transcription. SAGA (Spt-Ada-Gcn5 Acetyltransferase) is an evolutionary conserved coactivator complex. The core module scaffolds the entire SAGA complex and adopts a histone octamer-like structure, which consists of six histone fold domain (HFD)-containing proteins forming three histone fold (HF) pairs, to which the double HFD-containing SUPT3H adds an HF pair. Spt3, the yeast ortholog of SUPT3H, interacts genetically and biochemically with the TATA binding protein (TBP) and contributes to global RNA polymerase II (Pol II) transcription. Here we demonstrate that i) SAGA purified from human U2OS or mouse embryonic stem cells (mESC) can assemble without SUPT3H; ii) SUPT3H is not essential for mESC survival, iii) SUPT3H is required for mESC growth and self-renewal, and iv) the loss of SUPT3H from mammalian cells affects the transcription of only a specific subset of genes. Accordingly, in the absence of SUPT3H no major change in TBP accumulation at gene promoters was observed. Thus, SUPT3H is not required for the assembly of SAGA, TBP recruitment, or overall Pol II transcription, but plays a role in mESC growth and self-renewal. Our data further suggest that yeast and mammalian SAGA complexes contribute to transcription regulation by distinct mechanisms.

## Introduction

Formation of the transcription preinitiation complex (PIC), containing RNA polymerase II (Pol II) and six general transcription factors (GTFs), is a major regulatory step in eukaryotic gene expression (1,2). PIC formation is mediated by TFIID and the accessibility of the transcription machinery to template DNA is controlled by co-activator complexes. The SAGA complex (Spt-Ada-Gcn5 acetyltransferase) is an evolutionary conserved, multifunctional co-activator composed of about 18-20 subunits (3,4). Early genetic studies, predominantly performed in yeast, have established that SAGA is organized into distinct functional and structural entities, now called modules. These modules comprise a structural core, histone acetyltransferase (HAT), histone deubiquitinase (DUB), TBP-binding, and transcription factor (TF)-binding activities (5).

*Saccharomyces cerevisiae* (y) *SPT3* was isolated as an allele-specific suppressor of mutation in *TBP* coding for the TATA binding protein (TBP), suggesting that Spt3 may play a role in recruiting yeast SAGA (ySAGA) to promoters (6-8). The TBP-binding module of ySAGA comprises Spt3 and Spt8, which both interact with TBP directly (9,10). Recent high resolution cryo-electron microscopy (cryo-EM) structures of ySAGA complexes indicated that the core structural module contains a histone octamer like structure consisting of seven histone fold domain (HFD)-containing proteins, including Ada1/Taf12, Taf6/Taf9 and Taf10/Spt7, which form three HF pairs, and Spt3, which harbours two HF domains forming an intramolecular HF pair (10-12). Interestingly, Spt3 is homologous to the corresponding Taf11/Taf13 HF heterodimer in TFIID, and Spt3 of ySAGA binds the same side of TBP as does the Taf11/Taf13 HF pair (12-16). Purified recombinant TBP binds to the histone octamer-like structure of ySAGA through Spt3 and Spt8 (10,14), further arguing that yeast SAGA contributes to TBP delivery to core promoters. Indeed, structural and biochemical evidence suggest a model by which Spt3 prevents spurious TBP binding to DNA through steric hindrance, which is relieved by the synergistic binding of TFIIA and a cognate TATA element to TBP (10). In agreement, transcription of stress-inducible genes containing a TATA box in their promoters is more sensitive to SAGA subunit mutations, than to TFIID subunit mutations (17). However, recent analyses in yeast based on quantification of newly synthetized mRNA, demonstrated that both TFIID and SAGA are required for the transcription of almost all Pol II genes, although each subunit may act through different mechanisms and thus to different extent (18-20).

The modular organization of mammalian SAGA complexes is very similar to that of their yeast counterparts (4). For example, their histone octamer-like structure is built up by three homologous histone fold pairs, TADA1/TAF12, TAF6L/TAF9/9b, and TAF10/SUPT7L, and SUPT3H, the mammalian homologue of ySpt3 (21,22). However, metazoan SAGA complexes are lacking a Spt8 homologue (4), suggesting that TBP interactions with mammalian SAGA complexes may be different from that of yeast SAGA. In addition, fly and mammalian SAGA complexes contain also a splicing module (4,22,23).

Here we investigated the role of SUPT3H in SAGA complex assembly, Pol II transcription regulation and TBP recruitment to target gene promoters in human U2OS and mouse embryonic stem cells (ESCs). Our results indicate that, in both cell types, SAGA subunit composition is not affected by the absence of SUPT3H, similar to yeast (9,24,25). In contrast, we observed a striking divergence between ySpt3 and mammalian SUPT3H orthologues in their regulatory roles. While we previously showed that yeast Spt3 contributes to global Pol II transcription (18), we show here that SUPT3H is required only for the expression of a limited number of genes in both cell types. TBP recruitment experiments further show that absence of SUPT3H does not affect TBP binding to selected gene promoters. Despite these limited regulatory roles, we found that *Supt3* gene inactivation in mESCs affects growth and self-renewal. Together these data suggest that the role of SUPT3H in gene transcription diverged substantially between yeast and mammals, suggesting that mammalian SAGA complexes contribute to transcription regulation by distinct mechanism, or that SAGA-dependent TBP delivery to promoters requires other subunits.

## Materials and Methods

### Reagents

Reagents are described in Supplementary Table S1.

### Biological Resources

Biological resources are described in Supplementary Table S2, and see below:

#### a) Generation of stable Flag-SUPT3H overexpressing U2OS cell lines

The coding region (CDS) of the human *SUPT3H* gene was PCR amplified from the pREV-*SUPT3H* vector (26) using HA 13662980/1 primers described in Supplementary Table S3, and Phusion™ High Fidelity polymerase (Thermo Fisher Scientific, Cat# F-530), following manufacturer’s instruction. This first PCR product was purified on gel using NucleoSpin Gel and PCR Clean-up, Mini kit for gel extraction and PCR clean up (Machery-Nagel, Cat# 740609.50). A second round of PCR was performed to amplify this first obtained PCR product and eliminate pREV-*SUPT3H* matrix, using the same primers. Final product was purified on gel. This cDNA was cloned into pSG5-puro A frame expression vector (27). First the *SUPT3H* PCR product was digested with Xho I and Sma I type II restriction enzymes from New England Biolabs, and purified using Nucleospin^®^ PCR clean-up kit (Machery Nagel Cat# 740609.50). Then ligated into the XhoI and SmaI fragment of the pSG5-puro expression vector from an SV40 promoter, digested and purified same way as described above, using T4 DNA ligase (Biolabs, Cat# M0202). This product was transformed in competent DH5α and ampicillin resistant colonies were selected after. Final plasmid constructs were verified by sequencing using.

To obtain a U2OS cell line stably expressing the SUPT3H protein, U2OS cells were transfected with 5 *μ*g of circular pSG5puro-*hSUPT3H* plasmid, using Lipofectamine2000 following manufacturer’s instruction (ThermoFisher Scientific, Cat# 11668019). One single cell per well was seeded in 96-well plates using the BD Biosciences FACSAria II (BD Biosciences) apparatus. After a 10-days puromycin selection, positive cell clones were selected by western blot analysis using the anti-Flag and anti-SUPT3H antibody.

#### b) Generation of *Supt3*^-/-^ mutant mESC lines

Mouse E14 ESCs were transfected with a plasmid construct encoding for two sgRNAs (sequences in Supplementary Table S4) as well as a Cas9-GFP fusion protein at a confluency of 70-80% using Lipofectamine2000 (ThermoFisher Scientific, Cat# 11668019) following manufacturer’s instruction. Two days after transfection, cells were selected for expression of the Cas9-GFP fusion protein by fluorescence activated sorting (FACS). Five 96-well plates were seeded with one GFP positive cell per well using the BD FACSAriaTM II (BD Biosciences) apparatus. Clones were amplified, genomic DNA extracted and knockout clones were selected by PCR, using the Phire direct PCR kit (Thermo Scientific, Cat# F-1265) following manufacturer’s instructions. Primer sequences for PCR are shown in Supplementary Table S3. Homozygous mutant clones were sequenced using primers spanning the deletion site (Supplementary Table S3).

### Cell culture conditions

Human U2OS osteosarcoma cells (HTB-96; ATCC) and U2OS-Fl-SUPT3H cells were cultured using DMEM medium supplemented with 10% foetal calf serum (Sigma Aldrich, Cat# F7524) and 40 *μ*g/ml gentamycin (KALYS, Cat# G0124-25). Cells were grown at 37°C with 5% CO_2_ levels.

Mouse ES E14 cells were cultured on plates coated with 0.1% gelatin solution in 1x PBS (Dutcher, Cat# P06-20410) using DMEM medium supplemented with 15% foetal calf serum ES-tested (ThermoFisher Scientific, Cat# 10270-106), 2 mM ʟ-glutamine (ThermoFisher Scientific, Cat# 25030-024), 0.1% β-mercaptoethanol (ThermoFisher Scientific, Cat# 31350-010), 100 UI/ml penicillin and 100 μg/ml streptomycin (ThermoFisher Scientific, Cat# 15140-122), 0.1 mM non-essential amino acids (ThermoFisher Scientific, Cat# 11140-035) and 1,500 U/ml leukaemia inhibitory factor (home-made). For medium described as FCS+LIF+2i medium, 3 μM CHIR99021 (Axon Medchem, Cat# 1386) and 1 μM PD0325901 (Axon Medchem, Cat# 1408) were added freshly to the medium. Cells were grown at 37°C with 5% CO_2_ levels.

*Schizosaccharomyces pombe* cells were grown in autoclaved YES medium (yeast extract, adenine, histidine, uracil, leucine, lysine, 3% glucose) at 32°C.

### Clonal assays of mouse ESCs

For clonal assay analyses, 1500 to 3000 cells, which had been adapted to the respective media through at least three passages, were plated in wells of 6-well plates. Medium was changed every other day. On the sixth day, colonies were washed twice with 1x PBS before fixation with 4% Paraformaldehyde (Electron Microscopy Sciences, Cat# 15710, 16% solution) for 30 minutes followed by two washes with 1x PBS. To assess the alkaline phosphatase (AP) activity of mouse ESC colonies, Alkaline Phosphatase Kit (Vector Laboratories, Cat# SK-5100) was used following manufacturer’s instructions. Colonies were stained with AP for 5-10 minutes. For clonal assay analyses in FCS+LIF medium, an additional staining with crystal violet was performed after AP staining. Colonies were stained with 0.1% crystal violet solution for at least 30 minutes.

### Quantification of clonal assays

For clonal assay analyses in FCS+LIF+2i medium, colony areas were measured automatically using ImageJ software. For clonal assay analyses in FCS+LIF medium, the colonies were counted manually using the ImageJ interface. We considered colonies as AP+ colonies if they either stained entirely red or if they possessed a center of red cells surrounded by unstained cells. The total number of colonies as assessed by crystal violet staining was used for normalization between replicates.

### Metabolic labelling

Metabolic labelling of newly synthesized RNA was adapted from previously described protocols (28-30). In brief, the nucleoside analogue 4-thiouridine (4sU) (Glentham Life Sciences, Cat# GN6085) was added to the cell culture medium at a final concentration of 500 μM for a 20-minute pulse for mouse ES E14, human U2OS cells or Schneider S2 cells. After the labelling period, the medium containing 4sU was removed, the cells were washed with ice cold 1x PBS and immediately lysed using TRI^®^ Reagent (Molecular Research Center Inc., Cat# TR 188).

*S. pombe* cultures were grown to an OD600 of 0.8. 4-thiouracil (Sigma Aldrich, Cat# 440736) was freshly dissolved in DMSO and added to the cultures at a final concentration of 1 mM. Labelling was performed for 6 minutes. After this time period, yeast cells were pelleted, washed with ice-cold 1x PBS and aliquoted before being flash frozen and stored at -80°C.

### Total RNA extraction

Total RNA was extracted following TRI^®^ Reagent (Molecular Research Center Inc., Cat# TR 188) manufacturer’s instruction. To remove any potential genomic DNA contamination from the total RNA extracts, the TURBO DNA-*free*^™^ Kit (ThermoFisher Scientific, Cat# AM1907) was used following manufacturer’s instructions for rigorous DNase treatment.

For total RNA extraction of yeast cells, the RiboPure^™^ RNA Purification Kit (ThermoFisher Scientific, Cat# AM1926) was used following manufacturer’s instruction.

### Reverse Transcription

Reverse Transcription (RT) was performed with 2 μg total RNA and using 3.2 μg random hexamer primers (ThermoFisher Scientific, Cat# SO142) and Transcriptor Reverse Transcriptase (Roche, Cat# 03531287001) following manufacturer’s instruction.

### qPCR

For qPCR, the cDNA samples were amplified using LightCycler^®^ 480 SYBR^®^ Green 2x PCR Master Mix I (Roche, Cat# 04887352001) and 0.3 or 0.6 μM of forward and reverse primer respectively. The primer pairs used for qPCR are listed in Supplementary Table S3. The qPCR was conducted using a LightCycler^®^ 480 (Roche). For the assessment of mRNA levels, the obtained threshold-values were used to calculate the relative gene expression using the 2^-ΔΔCT^ method and considering the individual primer pair efficiencies (31). For TBP ChIP-qPCR, the percentage of input was calculated.

### Newly synthesized RNA purification

The purification of newly synthesized RNA was based on previously described protocols (28-30). As spike-in, labelled *S. pombe* total RNA was added to labelled mouse ESC total RNA preparations in a ratio 1:10; or labelled *S. pombe* total RNA was added to labelled U2OS total RNA preparations in a ratio of 1:25 respectively, to a final amount of 200 μg of total RNA prior to newly synthesized RNA purification. The RNA was precipitated and resuspended in 130 *μ*L RNase-free water (Sigma Aldrich, Cat# 95284) and sonicated using the following settings on a Covaris E220 instrument: 1 % duty factor, 100 W, 200 cycles per burst, 80 seconds. Fragment size ranged from 10 kb to 200 bp. For purification, the fragmented total RNA was incubated for 10 minutes at 60°C and immediately chilled on ice for 2 minutes to open secondary RNA structures. The 4sU-labelled RNA was thiol-specific biotinylated by addition of 200 μg EZ-link HPDP-biotin (ThermoFisher Scientific, Cat# 21341), biotinylation buffer (10 mM Hepes-KOH pH 7.5 and 1 mM EDTA) and 20% DMSO (Sigma Aldrich, Cat# D8418). Biotinylation was carried out for 3 hours at 24°C in the dark and with gentle agitations. After incubation, excess biotin was removed by adding an equal volume of chloroform and centrifugation at 16,000 g for 5 minutes at 4°C. RNA was precipitated from the aqueous phase by adding 0.1 volumes of 5 M NaCl and an equal volume of 100% isopropanol followed by centrifugation at 16,000 g for 30 minutes at 4°C. After washing with 75% ethanol the RNA pellet was resuspended in 100 *μ*L of RNase-free water and denatured for 10 minutes at 65°C followed by immediate chilling on ice for 5 minutes. The samples were incubated with 100 *μ*L of streptavidin-coated μMACS magnetic beads (Miltenyi Biotec, Cat# 130-074-101) for 90 minutes at 24°C under gentle agitations. The μMACS columns (Miltenyi Biotec, Cat# 130-074-101) were placed on a MACS MultiStand (Miltenyi Biotec) and equilibrated with washing buffer (100 mM Tris-HCl pH 7.5, 10 mM EDTA, 1 M NaCl, 0.1% Tween20) before applying the samples twice to the columns. The columns were then washed one time with 600 *μ*L, 700 *μ*L, 800 *μ*L, 900 *μ*L and 1 ml washing buffer before eluting the newly synthesized RNA with two washes of 100 *μ*L 0.1M DTT. The isolated newly synthesized RNA was recovered either using RNeasy MinElute Cleanup Kit (Qiagen, Cat# 74204) following manufacturer’s instruction or by precipitation. Libraries were prepared and sequenced with 1x 50 base pairs on a HiSeq4000 machine (Illumina).

### Library preparation of 4sU RNA-seq

For U2OS cells, 4sU RNA-Seq libraries were generated from 15 ng of purified, newly synthesized RNA using Illumina Stranded Total RNA Prep, Ligation with Ribo-Zero Plus kit and IDT for Illumina RNA UD Indexes, Ligation (Illumina, San Diego, USA, Cat# 20040525 and 20040553/4, respectively), according to manufacturer’s instructions. Briefly, abundant ribosomal RNAs were depleted by hybridization to specific DNA probes and enzymatic digestion. The depleted RNAs were purified and fragmented using divalent cations at 94°C for 2 minutes. After random hexamers annealing, fragmented RNAs were then reverse transcribed into first strand complementary DNA (cDNA). Second strand cDNA synthesis further generated blunt-ended double-stranded cDNA and incorporated dTTP in place of dUTP to achieve strand specificity by quenching the second strand during amplification. Following A-tailing of DNA fragments and ligation of pre-index anchors, PCR amplification was used to add indexes and primer sequences and to enrich DNA libraries (30 sec at 98°C; [10 sec at 98°C, 30 sec at 60°C, 30 sec at 72°C] x 12 cycles; 5 min at 72°C). Surplus PCR primers were further removed by purification using SPRIselect beads (Beckman-Coulter, Villepinte, France, Cat# B23319) and the libraries were sequenced with 1x 50 base pairs on a HiSeq4000 machine (Illumina).

For mESCs, 4sU RNA-seq libraries were generated from 15 to 50 ng of purified, newly synthesized RNA using TruSeq Stranded Total RNA LT Sample Prep Kit with Ribo-Zero Gold (Illumina, San Diego, CA, Cat# RS-122-2301) according to the Illumina protocol with the following modifications. 4sU-labelled RNA was cleaned up using 1.8X RNAClean XP beads and fragmented using divalent cations at 94oC for 1 minutes without depletion of rRNA. While, double stranded cDNA synthesis and adapter ligation were performed according to manufacturer instructions, the number of PCR cycles for library amplification was reduced to 10 cycles. After purification using SPRIselect beads (Beckman-Coulter, Villepinte, France, Cat# B23319), the libraries were sequenced with 1x 50 base pairs on a HiSeq4000 machine (Illumina).

### Data analysis of 4sU RNA-seq

Reads were preprocessed using CUTADAPT 1.10 (32) in order to remove adaptors and low-quality sequences and reads shorter than 40 bp were removed for further analysis. rRNA sequences were removed for further analysis. For mESC samples VQFR25, VQFR26, VQFR29, VQFR30, remaining reads were aligned to a hybrid genome composed of mm10 and ASM294v2 assemblies of *M. musculus* and *S. pombe* genomes respectively with STAR 2.5.3a (33). For samples VQFR188, VQFR189, VQFR191, VQFR192, the hybrid genome was composed of hg38 and ASM294v2 assemblies of *H. sapiens* and *S. pombe* genomes respectively. Gene quantification was performed with htseq-count 0.6.1p1 (34), using “union” mode and Ensembl 93 annotations for all organisms except for *S. pombe* where Ensembl Fungi 41 annotations were used. For 4sU-seq data, “type” option was set to “gene” in order to take also into account reads aligned onto introns. Differential gene expression analysis was performed using DESeq2 1.16.1 (35) Bioconductor R package on *H. sapiens* or *M. musculus* counts normalized with size factors computed by the median-of-ratios method proposed by Anders and Huber (36) based on the spike-in counts (using the following options: cooksCutoff=TRUE, independentFiltering=TRUE, alpha=0.05). P-values were adjusted for multiple testing using the Benjamini and Hochberg method (37). For subsequent data analyses and visualization, only protein-coding genes were considered. Further, a threshold of 100 or 50 reads was used to define expressed genes in the U2OS or mESC datasets respectively.

For TATA-less and TATA-box promoter analysis (violin plots in Fig. 2E and Fig. 5E), TATA-box containing promoters were extracted for the human (hg38) and mouse (mm10) genomes using the Eukaryotic Promoter Database (EPD, https://epd.epfl.ch//index.php, (38)).

### Whole cell extract preparation from U2OS cells

The required number of cells were trypsinized, transferred into 1.5 ml Eppendorf tubes, centrifuged 600 g 4°C for 2 min, and washed twice with 1ml 1x PBS. Pellets were resuspended in 1 packed cells volume (PCV) extraction buffer (400 mM KCl, 20 mM Tris-HCl pH 7.5, 20% glycerol, 2 mM DTT, 1x protease inhibitor cocktail). Tubes were frozen and thawed subsequently 4 times (from liquid nitrogen to ice) and centrifuged at 14,000 g 4°C for 10 min. The supernatant was stored at -80°C.

### Human cell nuclear extract preparation

To enrich extracts for nuclear proteins, cells were harvested and washed twice with 1x PBS. Cell pellets were resuspended in 4 times PCV of hypotonic buffer (50 mM Tris pH 7.9, 1 mM EDTA, 1 mM DTT and 1x protein inhibitor cocktail), left to swell for 30 minutes on ice, then dounced 10 times using a B dounce homogenizer to break cytoplasmic membrane. After a 10 minutes centrifugation at 1,000-1,800 g, 4°C, supernatant was removed and the pellet resuspended in a high salt buffer (50 mM Tris pH 7.9, 25 % glycerol, 500 mM NaCl, 0.5 mM EDTA, 1 mM DTT and 1x protein inhibitor cocktail). To break the nuclear membranes, suspension was homogenized by douncing 20 times using a B dounce, then incubated at 4°C (under stirring), for 30 minutes prior to centrifugation at 10,000 g for 20 minutes at 4°C. The supernatant was dialyzed o/n at 4°C against an isotonic salt buffer (50 mM Tris pH 7.9, 20 % glycerol, 5 mM MgCl_2_, 100 mM KCl, 1 mM DTT and 1x protein inhibitor cocktail). The dialyzed fraction was kept as nuclear extract.

### Mouse ESC nuclear extract preparation

To enrich extracts for nuclear proteins, cells were harvested and washed twice with 1x PBS. Cell pellet was resuspended in hypotonic buffer (10 mM Tris-HCl pH 8.0, 1.5 mM MgCl_2_, 10 mM KCl and 1x protein inhibitor cocktail) and dounced 10 to 20 times using a B dounce homogenizer to isolate the nuclei. After centrifugation at 10,000 g for 10 minutes at 4°C, supernatant was removed and pellet resuspended in high salt buffer (20 mM Tris-HCl pH 8.0, 25% glycerol, 1.5 mM MgCl_2_, 0.2 mM EDTA, 450 mM NaCl, 0.1% NP40 and 1x protein inhibitor cocktail). Suspension was homogenized by douncing as described before prior to centrifugation at 10,000 g for 10 minutes at 4°C. The supernatant was kept at -80°C as nuclear extract.

### Antibodies

The list of antibodies used in this study is shown in Supplementary Table S5. Anti-TADA2B (3122) polyclonal antibody (pAb) was obtained by immunization of rabbits with the C-terminal region (amino acids 221-420) of hTADA2B (Q86TJ2). For this, the *TADA2b* cDNA fragment was PCR amplified and cloned in pET15b vector (Novagen) using Nde I and Bam HI sites. The 6xHis-TADA2B protein fragment was expressed in *E. coli* (BL21), centrifuged, lysed in a buffer L, containing 50 mM Tris-HCl pH 7.5, 400 mM NaCl, 10% glycerol, 1 mM DTT and 1x protein inhibitor cocktail. Recombinant proteins were in inclusion bodies, which were solubilized in buffer L containing 8 M urea. Proteins were purified under denaturing conditions using a Ni^2+^-NTA column, eluted with 300 mM imidazole and dialyzed in 1x PBS. Rabbits were immunized with the purified proteins for 6 weeks as described in (39). The obtained crude rabbit sera were then purified on an Affigel column on which the purified protein fragment was immobilized. Then the column was extensively washed with 1x PBS, pAb3122 eluted with 0.1 M glycine pH 2.5 buffer and immediately neutralized with 2 M Tris pH 8.8.

### Immunoprecipitation protocol from human cell extracts

Prior to immunoprecipitation, protein-A sepharose and ANTI-FLAG^®^ M2 Affinity Gel beads (Sigma Aldrich, Cat# A2220) were washed three times with 1x PBS and two times with IP100 buffer (25 mM Tris-HCl 7.5, 5 mM MgCl_2_, 10 % glycerol, 0,1 % NP40, 100 mM KCl, 2 mM DTT and 1x protein inhibitor cocktail), prior to use. Nuclear extracts were pre-cleared with 1/10 volume of packed bead volume for 1 hours at 4°C with agitation. For antibody binding, packed bead volume corresponding to 1/10 volume of input extract was incubated with the respective antibodies for 1 hours at room temperature, with agitation. Protein-A antibody bound sepharose beads were washed twice with IP500 (25 mM Tris-HCl 7.5, 5 mM MgCl2, 10% glycerol, 0,1% NP40, 500 mM KCl, 2 mM DTT and 1x protein inhibitor cocktail) buffer and then with IP100 buffer. Pre-cleared extracts were then incubated overnight with washed protein-A antibody bound sepharose beads at 4°C. Protein complex bound beads were washed twice with IP500 buffer and twice with IP100 buffer. Complexes were subsequently eluted twice with one bead volume of 0.1 M glycine pH 2.8 buffer at room temperature and with agitation. Eluates were immediately neutralized to pH7.5 by adding the required quantity of Tris-HCl pH 8.8 buffer. Eluates were characterized by western blot or mass spectrometry analyses.

### Immunoprecipitation protocol for mESC nuclear extracts

Prior to immunoprecipitations, Protein-A or Protein-G Sepharose beads were washed three times with filtered 1x PBS and two times with IP100 buffer (25 mM Tris-HCl 7.5, 5 mM MgCl_2_, 10% glycerol, 0,1% NP40, 100 mM KCl, 2 mM DTT and 1x protein inhibitor cocktail). Nuclear extracts were pre-cleared with 1/5 of 50% bead slurry for 2 hours at 4°C with agitation. For antibody binding, the 50% bead slurry was incubated with 5-8 *μ*g of the respective antibodies for 2 hours at 4°C with agitation. After incubation, beads were washed three times with IP500 buffer and twice with IP100 buffer before addition of 1/5 volume of the 50% antibody-bead slurry to the pre-cleared nuclear extracts. Nuclear extracts were incubated with beads overnight at 4°C with agitation. After incubation, resins were washed three times with IP500 buffer and twice with IP100 buffer. Complexes were eluted from the beads using two subsequent 0.1 M glycine pH 2.8 elutions at room temperature and with agitation. Importantly, prior to anti-TAF10 IPs in mESC, nuclear extracts were depleted for TFIID by overnight incubation with beads coated with antibodies targeting the TFIID-specific subunit TAF7. This allowed to increase the purification efficiency for SAGA in anti-TAF10 IPs given that TAF10 is shared between the SAGA and TFIID complexes. All other IPs have been performed without pre-depletion. Eluates were then characterized by western blot or mass spectrometry analyses.

### Mass spectrometry

Protein mixtures were Trichloroacetic acid (Sigma Aldrich, Cat# T0699) - precipitated overnight at 4°C. Samples were then centrifuged at 14000 rpm for 30 minutes at 4°C. Pellet were washed twice with 1 ml cold acetone and centrifuged at 16000 g rpm for 10 minutes at 4°C. Washed pellet were then urea-denatured with 8 M urea (Sigma Aldrich, Cat# U0631) in Tris-HCl 0.1 mM, reduced with 5 mM TCEP (tris(2-carboxyethyl)phosphine) for 30 minutes, and then alkylated with 10 mM iodoacetamide (Sigma Aldrich, Cat# I1149) for 30 minutes in the dark. Both reduction and alkylation were performed at room temperature and under agitation (850 rpm). Double digestion was performed with endoproteinase Lys-C (Wako, Cat# 125-05061) at a ratio 1/100 (enzyme/proteins) in 8 M urea for 4h, followed by an overnight modified trypsin digestion (Promega, Cat# V5111) at a ratio 1/100 (enzyme/proteins) in 2 M urea. Both Lys-C and Trypsin digestions were performed at 37°C. Peptide mixtures were then desalted on C18 spin-column and dried on Speed-Vacuum before LC-MS/MS analysis. Samples were analyzed using an Ultimate 3000 nano-RSLC (Thermo Scientific, San Jose California) coupled in line with a LTQ-Orbitrap ELITE mass spectrometer via a nano-electrospray ionization source (Thermo Scientific, San Jose California). Peptide mixtures were loaded on a C18 Acclaim PepMap100 trap-column (75 *μ*m ID x 2 cm, 3 *μ*m, 100Å, Thermo Fisher Scientific) for 3.5 minutes at 5 *μ*L/min with 2% Acetonitrile MS grade (Sigma Aldrich, Cat# 1207802), 0.1% formic acid (FA, Sigma Aldrich, Cat# 94318)in H_2_O and then separated on a C18 Accucore nano-column (75 *μ*m ID x 50 cm, 2.6 *μ*m, 150Å, Thermo Fisher Scientific) with a 90 minutes linear gradient from 5% to 35% buffer B (A: 0.1% FA in H_2_O / B: 99% Acetonitrile MS grade, 0.1% FA in H_2_O), then a 20 minutes linear gradient from 35% to 80% buffer B, followed with 5 min at 99% B and 5 minutes of regeneration at 5% B. The total duration was set to 120 minutes at a flow rate of 220 nL/min. The oven temperature was kept constant at 38°C. The mass spectrometer was operated in positive ionization mode, in data-dependent mode with survey scans from m/z 350-1500 acquired in the Orbitrap at a resolution of 120,000 at m/z 400. The 20 most intense peaks (TOP20) from survey scans were selected for further fragmentation in the Linear Ion Trap with an isolation window of 2.0 Da and were fragmented by CID with normalized collision energy of 35%. Unassigned and single charged states were rejected. The Ion Target Value for the survey scans (in the Orbitrap) and the MS2 mode (in the Linear Ion Trap) were set to 1E6 and 5E3 respectively and the maximum injection time was set to 100 ms for both scan modes. Dynamic exclusion was used. Exclusion duration was set to 20 s, repeat count was set to 1 and exclusion mass width was ± 10 ppm. Proteins were identified by database searching using SequestHT (Thermo Fisher Scientific) with Proteome Discoverer 2.4 software (PD2.4, Thermo Fisher Scientific) on *Homo sapiens* database (UniProt, reviewed, release 2020_11_27, 20309 entries) and *Mus musculus* database (UniProt, non-reviewed, release 2020_07_13, 55428 entries). Precursor and fragment mass tolerances were set at 10 ppm and 0.6 Da respectively, and up to 2 missed cleavages were allowed. Oxidation (M) was set as variable modification, and Carbamidomethylation (C) as fixed modification. Peptides were filtered with a false discovery rate (FDR) at 1%, rank 1 and proteins were identified with 1 unique peptide. Normalized spectral abundance factors (NSAF) were calculated for each protein as described earlier (40,41). First to obtain spectral abundance factors (SAF), spectral counts identifying a protein were divided by the protein length represented by the number of amino acids. Then to calculate NSAF values, the SAF values of each protein were divided by the sum of SAF values of all detected proteins.

### Anti-TBP chromatin immunoprecipitation (ChIP)

Cells were washed with 1x PBS, fixed at room temperature with 1% paraformaldehyde in 1x PBS for 20 minutes. Fixation was stopped by adding glycine to a final concentration of 125 mM for 10 minutes at room temperature. Cells were washed twice with ice-cold 1x PBS and collected by scrapping. Cells were centrifuged at 2000 g for 5 minutes at 4°C and washed once with ice-cold 1x PBS. Cells were lysed in L1 buffer (50 mM Tris-HCl pH 8.0, 2 mM EDTA, 0.1% NP40, 10% glycerol and protease inhibitory cocktail) and incubated on ice for 10 minutes before centrifugation at 2000g for 10 minutes at 4°C. Nuclei were resuspended in L2 buffer (0.5% SDS, 10 mM EDTA ph 8.0, 50 mM Tris-HCl pH 8.0 and protease inhibitory cocktail) and sonicated using a Covaris E210 sonicator to on average 300 bp fragments. Protein-G sepharose beads were washed twice with TE buffer (10 mM Tris-HCl pH 7.5, 1 mM EDTA) and blocked for three hours with 1 *μ*g/*μ*L denatured yeast tRNA and 1,5% fish-skin gelatine. Beads were washed twice with TE buffer and stored at 4°C. For ChIP, 50 *μ*g of chromatin was diluted with chromatin dilution buffer (0.01% SDS, 1.1% Triton-X 100, 1.2 mM EDTA, 167 mM NaCl, 16.7 mM Tris-HCl pH 8.0 and protease inhibitory cocktail) to a volume of 800 *μ*L and incubated with 30 *μ*L of blocked bead slurry for 1 hour at 4°C for preclearing. Five *μ*g of anti-TBP antibody (Abcam, cat # ab51841) was added to precleared chromatin and incubated overnight at 4°C. 50 *μ*L of bead slurry was added to samples and incubated for two to four hours at 4°C with overhead shaking. After centrifugation for 2 minutes at 1000 g, beads were washed two times with Low salt washing buffer (0.1% SDS, 1% Triton-X 100, 2 mM EDTA, 150 mM NaCl, 20 mM Tris HCl pH 8.0 and protease inhibitory cocktail) for 10 minutes, two times with high salt washing buffer (0.1% SDS, 1% Triton-X 100, 2 mM EDTA, 500 mM NaCl, 20 mM Tris HCl pH 8.0 and protease inhibitory cocktail) for 10 minutes, two times with LiCl washing buffer (250 mM LiCl, 1% NP40, 1% sodium deoxycholate, 1 mM EDTA, 10 mM Tris-HCl pH 8.0 and protease inhibitory cocktail) for 10 minutes and two times with TE buffer for 10 minutes. Bound fragments were eluted using with freshly prepared elution buffer (1% SDS and 0.1 M sodium bicarbonate) at room temperature. Chromatin was reverse crosslinked by addition of 0.2 M NaCl and 50 *μ*g/mL RNase A. Samples were incubated at 37°C for one hour. 20 *μ*g of Proteinase K was added and samples were incubated at 65°C overnight in a thermomixer. DNA was isolated by Phenol-chloroform purification and resuspended in water.

### Statistical analysis

All statistical tests were performed using either R (version 3.6.0). Two-sided Welch’s t-test were performed to compare colony areas after quantification with ImageJ between WT and mutant mESCs. Two-sided Wilcoxon rank-sum test was used to compare two means in all the other indicated comparisons. The statistical details for individual experiments can be found in the figure legends or Results section. This includes number of replicates and statistical tests performed. The statistical threshold was defined with a *p*-value of less than 0.05.

### Data availability

The 4sU-seq data discussed in this publication have been deposited in NCBI’s Gene Expression Omnibus (42) and are accessible through GEO Series accession numbers GSE175901. The mass spectrometry proteomics data have been deposited to the ProteomeXchange Consortium via the PRIDE (43) partner repository with the dataset identifier PXD026991.

## Results

### Human U2OS cells do not express the SUPT3H subunit of the SAGA complex

When analyzing RNA-sequencing data from the Human Protein Atlas (HPA) for mRNAs expressing different SAGA subunits in a panel of 64 human cell lines, we noticed that *SUPT3H* is not expressed in U2OS cells, a human osteosarcoma cell line, unlike the majority of other cell lines tested (Supplementary Figure S1A). To verify this observation, we designed primer pairs to amplify the whole coding sequence (CDS) of *SUPT3H* from cDNA prepared from U2OS and HeLa cells (Figure 1A, Supplementary Figure S1B). As an additional control, we generated an U2OS cell line stably expressing Flag-tagged wild type (WT) SUPT3H, which we refer to as U2OS-Fl-SUPT3H. In agreement with the HPA data, we could not detect the *SUPT3H* mRNA in U2OS cells, while it was clearly detectable in HeLa and U2OS-Fl-SUPT3H cells. Similar results were obtained when amplifying smaller fragments of the *SUPT3H* mRNA (Supplementary Figure S1D, scheme of primer positions Supplementary Figure S1B). Together these data indicate that the human U2OS osteosarcoma cell line does not express the SAGA subunit SUPT3H. Importantly, we could not detect major differences in the promoter sequence of the *SUPT3H* gene when comparing U2OS and HeLa cells, suggesting that the loss of *SUPT3H* expression is not caused by changes in the tested promoter sequence (Suppl. Fig. S1E). In addition, growth curve analyses revealed that U2OS-Fl-SUPT3H cells proliferate at a comparable rate than the parental U2OS cells (Supplementary Figure S1C).

**Figure 1:**
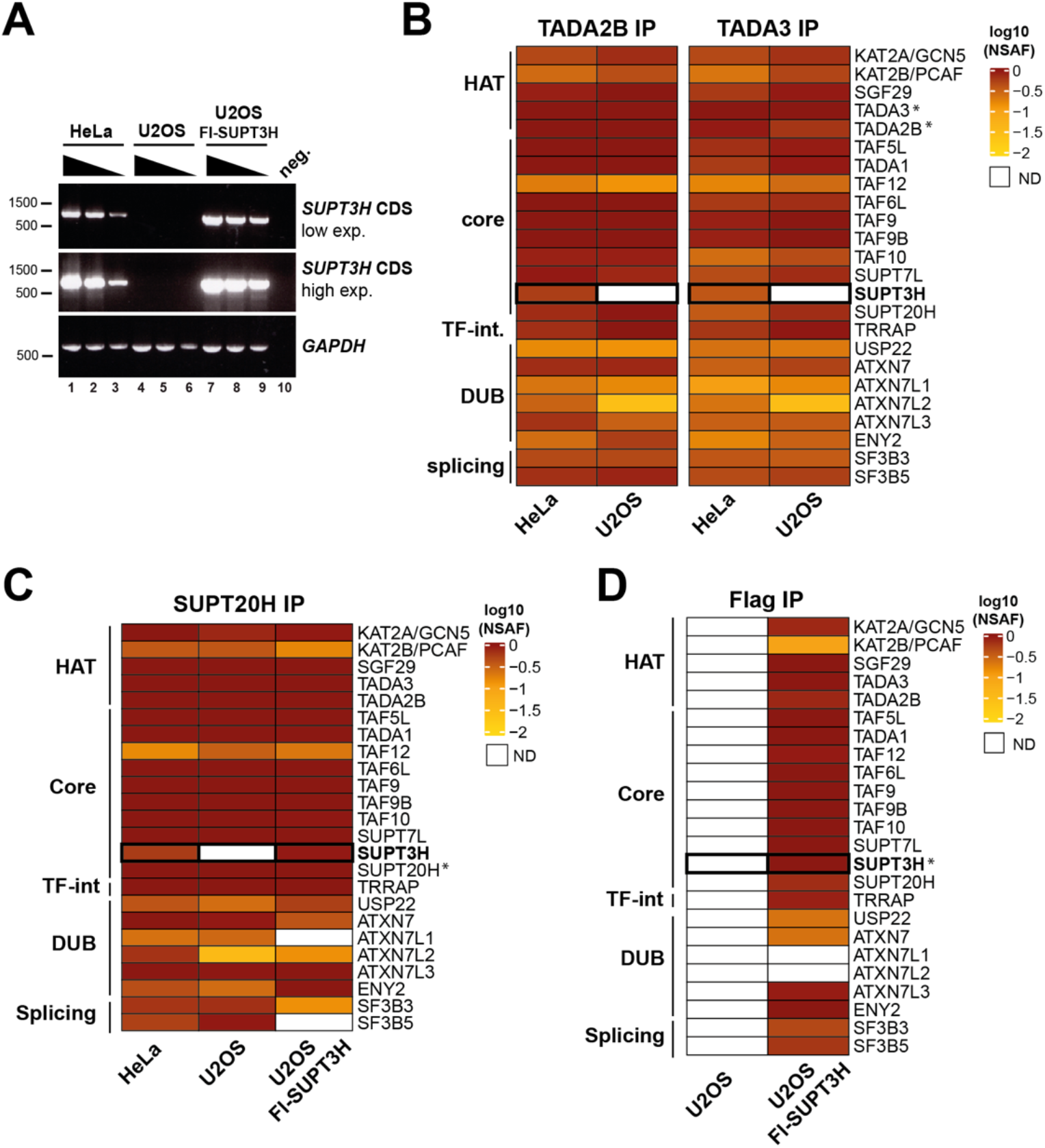
Absence of the SAGA subunit SUPT3H in human U2OS cells does not hinder formation of the SAGA complex. **(A)** PCR of the coding sequence (CDS) of *SUPT3H* using cDNA obtained from HeLa cells, U2OS cells or U2OS cells re-expressing Flag-SUPT3H (Fl-SUPT3H). Low and high exposures (exp.) are shown. *Gapdh* expression serves as loading control. n = 3 technical replicates. **(B)** Log10-transformed bait-normalized NSAF (Normalized Spectral Abundance Factor) values of mass spectrometry results from anti-TADA2B and anti-TADA3 immunoprecipitations (IP) of the SAGA complex from HeLa and U2OS nuclear extracts. n = 3 technical replicates in each IPs. **(C)** Log10-transformed bait-normalized NSAF values of mass spectrometry results from SUPT20H immunoprecipitations (IP) of the SAGA complex from HeLa, U2OS and U2OS Fl-SUPT3H nuclear extracts. n = 3 technical replicates. **(D)** Log10-transformed bait-normalized NSAF values of mass spectrometry results from Flag IPs shown in (Supplementary Figure S1F). n = 3 technical replicates. In (B, C, D) star (*) indicates the bait proteins. The distinct SAGA modules are indicated as HAT = histone acetyltransferase; TF-int = transcription factor-interacting; DUB = deubiquitylation. ND = not detected.

**Figure 2:**
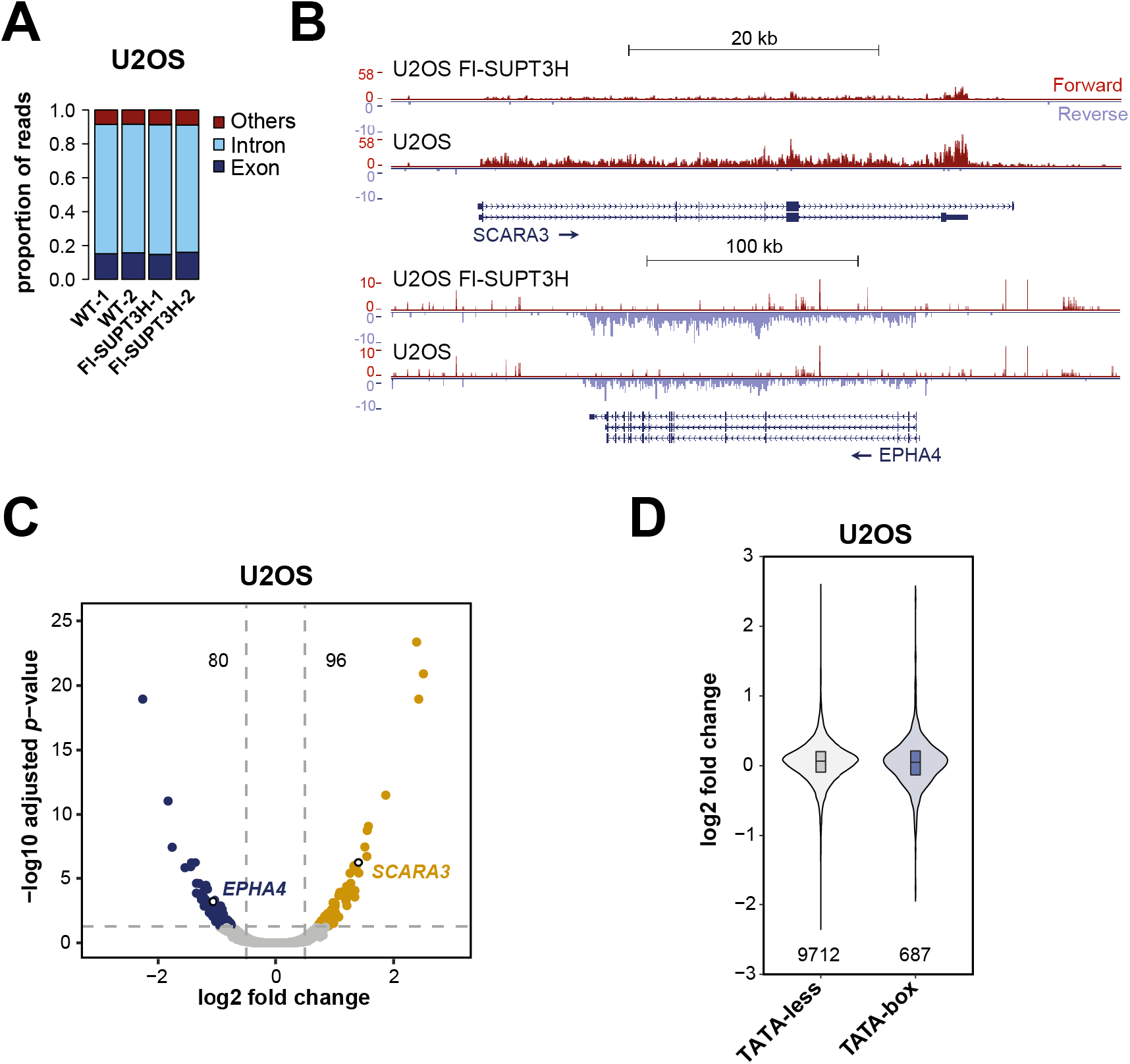
Only a small subset of Pol II-transcribed genes is responsive to the re-expression of SUPT3H in human U2OS cells. **(A)** Proportion of reads per genomic element for 4sU-seq experiments for two independent U2OS and Fl-SUPT3H clones. Exon = reads aligning to exons; Intron = reads aligning to introns, Others = reads aligning to intergenic regions, exon-intergenic junctions and exon-intron junctions. **(B)** UCSC genome browser views of 4sU-seq experiments on two differentially expressed genes (*SCARA3* and *EPHA4*) between U2OS Fl-SUPT3H and U2OS cells. Forward and reverse strands are shown. Arrows indicate direction of transcription. **(C)** Volcano plot representation of 4sU-seq with significantly down- or upregulated genes shown in dark blue and dark yellow respectively with numbers indicated on the top, using two independent U2OS and Fl-SUPT3H clones. A threshold of 100 reads was used to define expressed genes. The position of the two genes shown in (B) are indicated by circles. **(D)** Violin plot representation comparing the distribution of the log2 fold changes between TATA-less and TATA-box containing gene classes. The number of genes per category are indicated at the bottom. The statistical test performed is two-sided Welch’s t-test. Statistical significance (*p*<0.05) was not reached.

### Absence of SUPT3H does not affect human SAGA complex composition

As SUPT3H is a subunit of the core module of the SAGA complex interacting with several subunits of the complex, we were wondering if, and to which extent, SAGA is assembled in U2OS osteosarcoma cells lacking SUPT3H. To this end we prepared nuclear extracts from U2OS cells and carried out immunoprecipitation (IP) coupled to mass spectrometry using antibodies raised against SUPT20H (a core module subunit), TADA2B and TADA3 (two HAT module subunits). As a positive control we used human HeLa cell nuclear extracts (NEs) for IPs with the same antibodies. Quantitative mass spectrometry analyses of anti-SUPT20H, -TADA2B and -TADA3 affinity purifications showed that the three antibodies purified all SAGA subunits except SUPT3H from U2OS NEs, and all SAGA subunits from HeLa NEs (Figure 1B and 1C). Thus, the absence of SUPT3H does not affect the overall subunit composition of human SAGA. Importantly, anti-SUPT20H and anti-Flag IPs performed using nuclear extracts from U2OS-Fl-SUPT3H cells revealed that SUPT3H, when exogenously expressed, can incorporate in the SAGA complex in the U2OS nucleus (Figure 1C, 1D and Supplementary Figure S1F), indicating that the other subunits of the module are able to incorporate SUPT3H. To conclude, the absence of SUPT3H expression in U2OS cells allowed us to demonstrate that this subunit is dispensable for SAGA assembly.

### Restoring SUPT3H expression modifies RNA polymerase II transcription at a subset of genes in U2OS cells

The ability of SUPT3H to reconstitute an intact SAGA complex in U2OS-Fl-SUPT3H allowed us to test the role of SUPT3H in Pol II transcription in human cells. For this, we performed 4-thiouridine (4sU) labelling of nascent RNA coupled with sequencing of the labelled RNA (4sU-seq) (28-30), comparing two independent U2OS-Fl-SUPT3H clones with parental U2OS control cells (Supplementary Figure 2A). We observed a very similar enrichment in intronic reads in all samples (Figure 2A), confirming a reproducible enrichment of newly synthesized RNAs genome-wide. Differential expression analyses revealed that restoring SUPT3H expression has only a limited effect on nascent transcript levels (Figure 2C, 2D, Supplementary Figure S2B). Specifically, 176 genes were significantly deregulated in WT U2OS cells lacking SUPT3H compared to U2OS-Fl-SUPT3H cells at a threshold of 0.5 absolute log2 fold change and an adjusted *p*-value of 0.05, including 80 genes downregulated and 96 genes upregulated (Figure 2C, Supplementary Figure S2B). Only 33 and 37 genes are down-and upregulated, respectively, more than two-fold. Overall, no GO categories were significantly enriched amongst genes deregulated by *SUPT3H* overexpression and only very few genes are responsive to the re-expression of SUPT3H in U2OS cells.

Earlier studies suggested that ySAGA predominantly regulates TATA-box containing genes, presumably through Spt3 (7,44,45). We therefore compared expression changes between U2OS and U2OS-Fl-SUPT3H for genes with either a TATA-less or a TATA-box in their promoters, as defined by the Eukaryotic Promoter Database (Material and Methods) (Figure 2D). We observed no difference between the two gene classes, suggesting that TATA-box containing gene promoters do not show a stronger sensitivity to SUPT3H re-expression than TATA-less promoters in human U2OS cells. In conclusion, SUPT3H appears to regulate the transcription of only a few genes in osteosarcoma U2OS cells, independently of the presence a TATA-box element in their promoter. This observation is in marked contrast with the strong, global decrease of nascent transcript levels observed upon deletion of *SPT3* in *S. cerevisiae*.

### Loss of mouse *Supt3* does not affect SAGA subunit composition in mouse embryonic stem cells

Our results so far indicate that the function of ySpt3 in RNA Pol II transcription has diverged substantially between yeast and humans. To explore this further, we next turned to non-cancerous diploid mammalian cells and used CRISPR/Cas9 genome editing to inactivate the *Supt3* gene in mouse E14 ESCs. We obtained two individual clones with a homozygous deletion of exon 3 from the *Supt3* gene (Figure 3A), resulting in an out of frame stop codon. RT-qPCR analyses confirmed the deletion of the exon and the resulting decrease of remaining *Supt3* mRNA, presumably through non-sense decay (Figure 3B). Western blot analyses confirmed the deletion as SUPT3H is not detectable in extracts from knock-out mESCs (Figure 3C). To determine the role of SUPT3H on mouse SAGA assembly, we purified SAGA using anti-SUPT20H or anti-TAF10 antibodies from WT and *Supt3*^-/-^ ESCs nuclear-enriched cell extracts and analyzed their composition by mass spectrometry and western blot analyses. Both immune-purifications coupled to mass spectrometry analyses from *Supt3*^*-/-*^ cells identified all core, HAT and TF-binding SAGA subunits, with the exception of SUPT3H, indicating that mouse SAGA can also assemble without SUPT3H (Figure 3D). We note that the association of the DUB complex with SAGA was weaker in both anti-SUPT20H or anti-TAF10 IPs, but independently of the genotype of mESCs. These results confirm that, as observed in human U2OS cells, mouse SUPT3H has no major role in SAGA integrity in ESCs. Thus, SAGA central core can assemble without the intramolecular HF pair of SUPT3H, suggesting the SAGA can assemble around an hexameric core structure.

**Figure 3:**
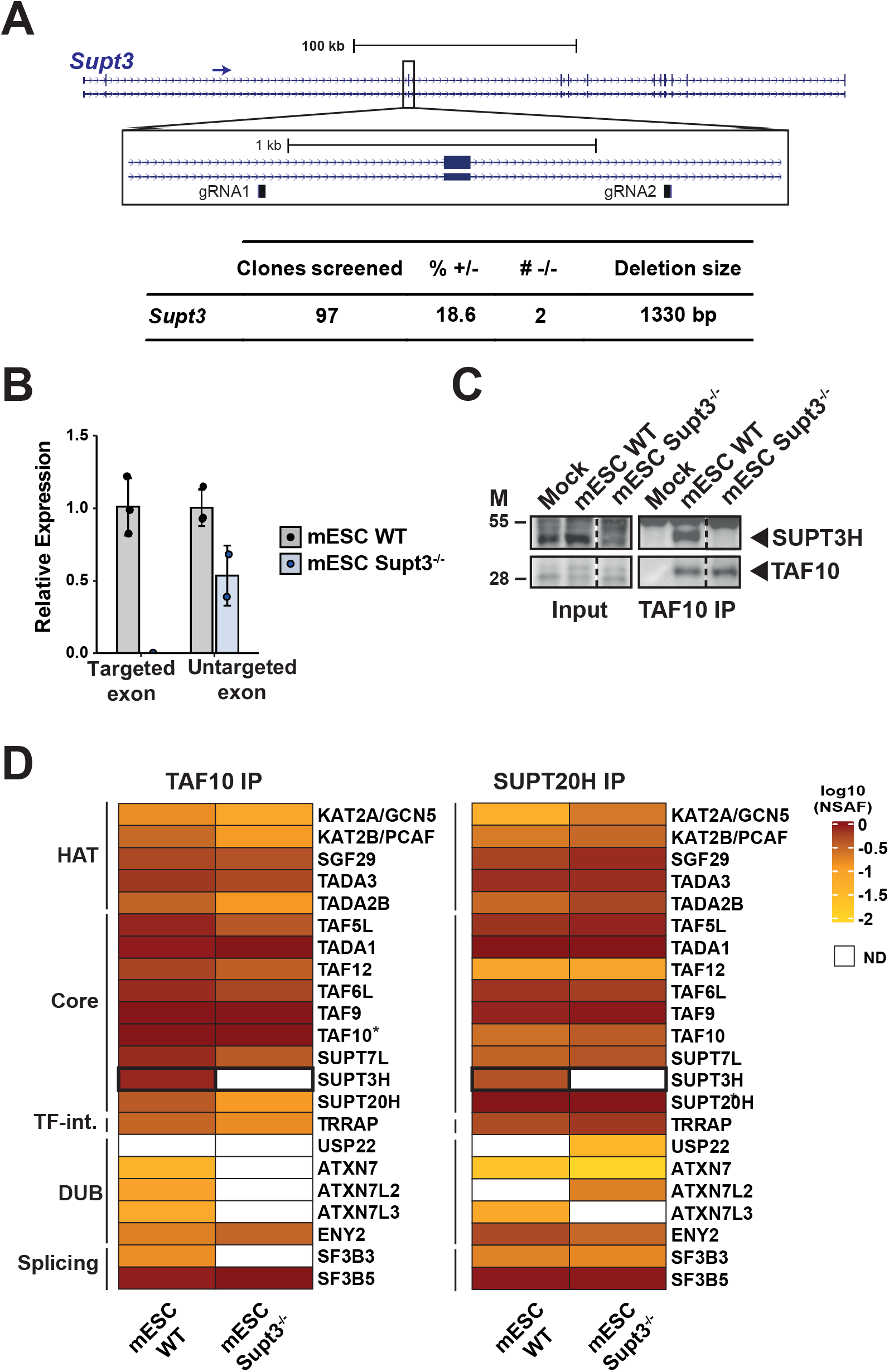
Loss of *Supt3* does not impair mouse ESC survival or formation of the SAGA complex. **(A)** Top, schematic representation of the mouse *Supt3* locus. The insert shows the position of the two gRNAs used to generate the *Supt3*^*-/-*^ cell lines. Bottom, table showing the number of clones screened, the percentage of heterozygous clones (+/-) and the number of homozygous (-/-) clones obtained as well as the deletion size. **(B)** Validation of *Supt3*^*-/-*^ cell lines by RT-qPCR revealing the absence of the targeted, out-of-frame exon and overall reduced levels of the exon-deleted *Supt3* mRNA. Error bars show mean ± standard deviation (SD) of two independent clones, with each assessment of the cell lines being based on the mean of three technical RT-qPCR replicates. **(C)** Western blot analysis of input and elution fractions of TAF10 immunoprecipitations (IP) from wildtype (WT mESCs) and *Supt3*^*-/-*^ mESCs. M = molecular weight markers (in kDa). **(D)** Log10-transformed bait-normalized NSAF values of mass spectrometry results from TAF10 IPs shown in (C) and SUPT20H IPs. n = three technical replicates each. Star (*) indicates the bait proteins. The distinct SAGA modules are indicated as HAT = histone acetyltransferase; TF-int = transcription factor-interacting; DUB = deubiquitylation. ND = not detected.

### *Supt3* is required for mouse ESC growth and self-renewal

The obtention of two independent *Supt3*^*-/-*^ mouse ESC lines indicates that *Supt3* is not essential for mouse ESC viability when cultured in medium containing foetal calf serum (FCS), leukemia inhibitory factor (LIF) and two small molecules that maintain efficient mouse ESC self-renewal (hereafter called FCS+LIF+2i medium). However, we observed in clonal assays that *Supt3*^*-/-*^ cells formed significantly smaller colonies compared to wildtype cells revealing that SUPT3H loss impairs mouse ESC growth (Figure 4A). In agreement, growth curve analyses revealed decreased proliferation of *Supt3*^*-/-*^ cells as compared to wildtype ESCs (Figure 4B).

**Figure 4:**
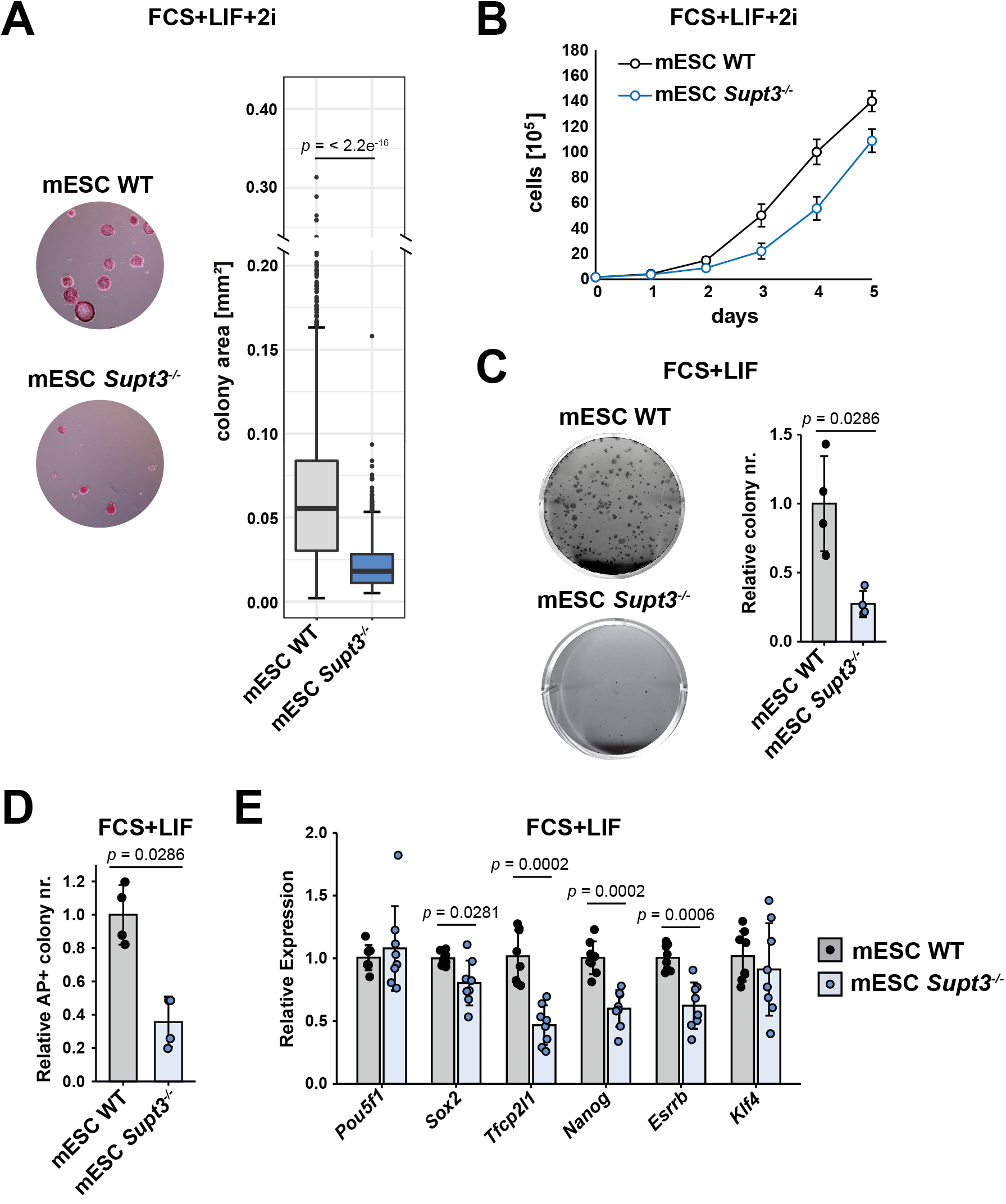
SUPT3H is required for mouse ESC growth and efficient self-renewal. **(A)** Clonal assays in FCS+LIF+2i medium comparing WT and *Supt3*^*-/-*^ mouse ESCs. Colonies were stained with alkaline phosphatase. Left, representative images; Right, quantification of colony areas using ImageJ. The statistical test performed is two-sided Welch’s t-test. Error bars show mean ± standard deviation (SD) of four replicates from two biological replicates, with two independent clones each. p value is indicated on the graph. **(B)** Growth curve analysis of viable cells comparing WT and *Supt3*^*-/-*^ mouse ESCs grown in FCS+LIF+2i medium for five days. Error bars show mean ± standard deviation (SD) of two biological replicates using two independent clones. **(C)** Clonal assays in FCS+LIF medium comparing WT and *Supt3*^*-/-*^ cells. Colonies were stained with crystal violet. Left, representative images; Right, quantification of colony numbers relative to WT cells. The statistical test performed is two-sided Wilcoxon rank-sum test. Error bars show mean ± standard deviation (SD) of four replicates from two biological replicates, with two independent clones each. p value is indicated on the graph. **(D)** Quantification of the number of colonies staining positive for alkaline phosphatase (AP+) in FCS+LIF medium normalized relative to WT cells. The statistical test performed is two-sided Wilcoxon rank-sum test. Error bars show mean ± standard deviation (SD) of four replicates from two biological replicates, with two independent clones each. p value is indicated on the graph. **(E)** Relative mRNA levels of pluripotency transcription factors (as indicated) comparing WT and *Supt3*^*-/-*^ mouse ESCs grown in FCS+LIF medium. Expression was normalized to RNA polymerase III transcribed genes (*Rn7sk* and *Rpph1*) as well as to WT cells. Error bars show mean ± standard deviation (SD) of at least 7 biological replicates (mean of three technical RT-qPCR replicates) using two independent clones each. Statistical test performed is two-sided Wilcoxon rank-sum test. Only statistically significant p values (*p*<0.05) are indicated.

**Figure 5:**
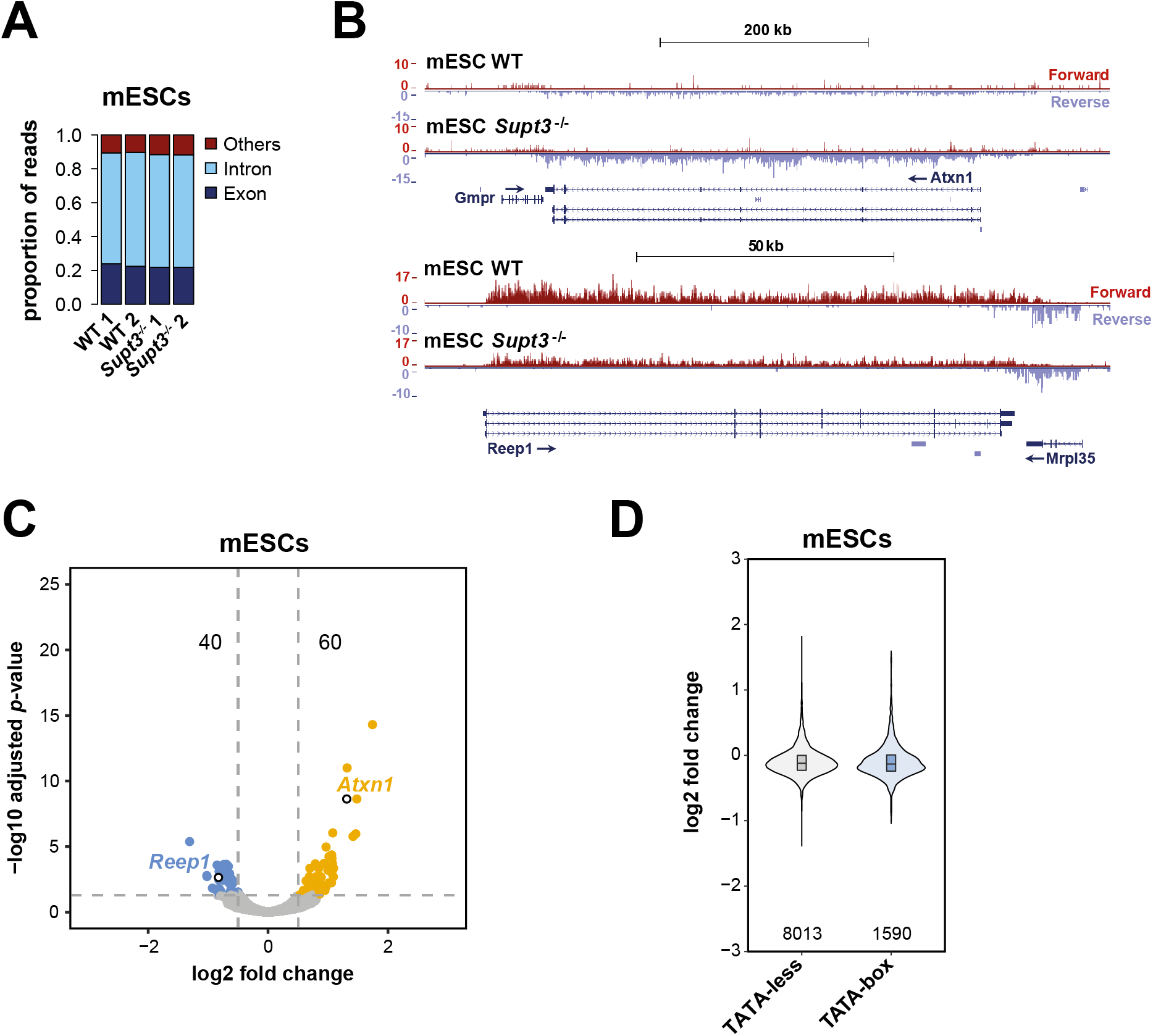
Loss of *Supt3* in mouse ESC has no major effect on Pol II transcription regulation in FCS+LIF+2i medium. **(A)** Proportion of reads per genomic element for 4sU-seq experiments in two independent wildtype (WT) and *Supt3*^*-/-*^ mouse ESC clones. Exon = reads aligning to exons; Intron = reads aligning to introns, Others = reads aligning to intergenic regions, exon-intergenic junctions and exon-intron junctions. **(B)** UCSC genome browser views of 4sU-seq experiments on two differentially expressed genes (*Atxn1* and *Reep1*) between WT and *Supt3*^*-/-*^ mouse ESCs. Forward and reverse strands are shown. Arrows indicate direction of transcription. **(C)** Volcano plot representation of 4sU-seq experiments with significantly down- or upregulated genes shown in light blue or yellow respectively with numbers indicated on the top, in two independent wildtype (WT) and *Supt3*^*-/-*^ mouse ESC clones. A threshold of 50 reads was used to define expressed genes. The position of the two differentially expressed genes shown in (B) are indicated by white circles with black border. **(D)** Violin plot representation comparing the distribution of the log2 fold changes between TATA-less and TATA-box containing genes. The number of genes per category are indicated at the bottom. The statistical test performed is two-sided Welch’s t-test.

To assess the importance of SUPT3H for the self-renewal capacities of ESCs, we performed clonal assays in medium without 2i (hereafter referred to as FCS+LIF medium) and observed about a 4-fold decrease in colony numbers in *Supt3*^-/-^ ESCs grown in FCS+LIF medium, as compared to WT cells (Figure 4C). As a read-out of the self-renewal capacities of ESCs, we measured the proportion of alkaline phosphatase positive (AP+) colonies. When compared to WT ESCs, the *Supt3*^*-/-*^ cells produced about 3-fold less AP+ colonies (Figure 4D), demonstrating that SUPT3H loss has a major impact on the self-renewal capacities of mouse ESCs.

### The loss of SUPT3H in mouse ESCs has no major effect on Pol II transcription

To determine the impact of SAGA lacking SUPT3H on Pol II transcription in mouse ESCs, we analyzed nascent RNA by 4sU-seq in FCS+LIF+2i medium in which *Supt3*^*-/-*^ ESCs can maintain their self-renewal capacities (as assessed by AP staining, Figure 4A). We could observe a similar enrichment of intronic reads in all samples (Fig. 5A). We observed that only a few genes, about 100, are differentially expressed between *Supt3*^*-/-*^ and WT ESCs (Fig. 5B, 5C and Supplementary Figure S2B). Specifically, the newly synthesized levels of about 40 transcripts were significantly decreased when applying a threshold of 0.5 log2 fold change and an adjusted *p*-value of 0.05. Conversely, the nascent levels of 60 transcripts increased in *Supt3*^-/-^ cells using the same thresholds (Supplementary Table 6). We could not find specific GO categories to be affected among the differentially expressed genes, however, the deregulation of some of these genes may explain the growth and self-renewal defects observed in *Supt3*^*-/-*^ mESCs.

Finally, we compared transcription changes between genes possessing TATA-less and TATA-box containing promoters (Fig. 5D). We observed no difference between the two gene classes suggesting that TATA-box containing promoters do not show a stronger sensitivity to the loss of *Supt3* than TATA-less promoters in mouse ESCs (Fig. 2D). These data together suggest that mouse SAGA complexes lacking Supt3, as observed in human U2OS cells, affect the expression of a small subset of genes in FCS+LIF+2i medium.

### Loss of SUPT3H in human U2OS cells or mouse ESCs does not significantly affect TBP binding

Genetic, biochemical and cryo-EM studies of ySAGA indicate that the Spt3 and Spt8 subunits bind to TBP and are involved in delivering TBP at specific promoters (6,7,10,11,46,47). We therefore wanted to assess whether SUPT3H has a role in recruiting TBP to mammalian gene promoters. Therefore, we performed TBP chromatin immunoprecipitation (ChIP)-coupled-qPCR experiments at selected gene promoters, in both U2OS and mESC cell lines (Figure 6A and 6B). We observed no difference in TBP occupancy at the tested gene promoters when comparing cells with and without SUPT3H. This suggests that TBP binding at these promoters does not require SUPT3H in human and mouse cells.

**Figure 6:**
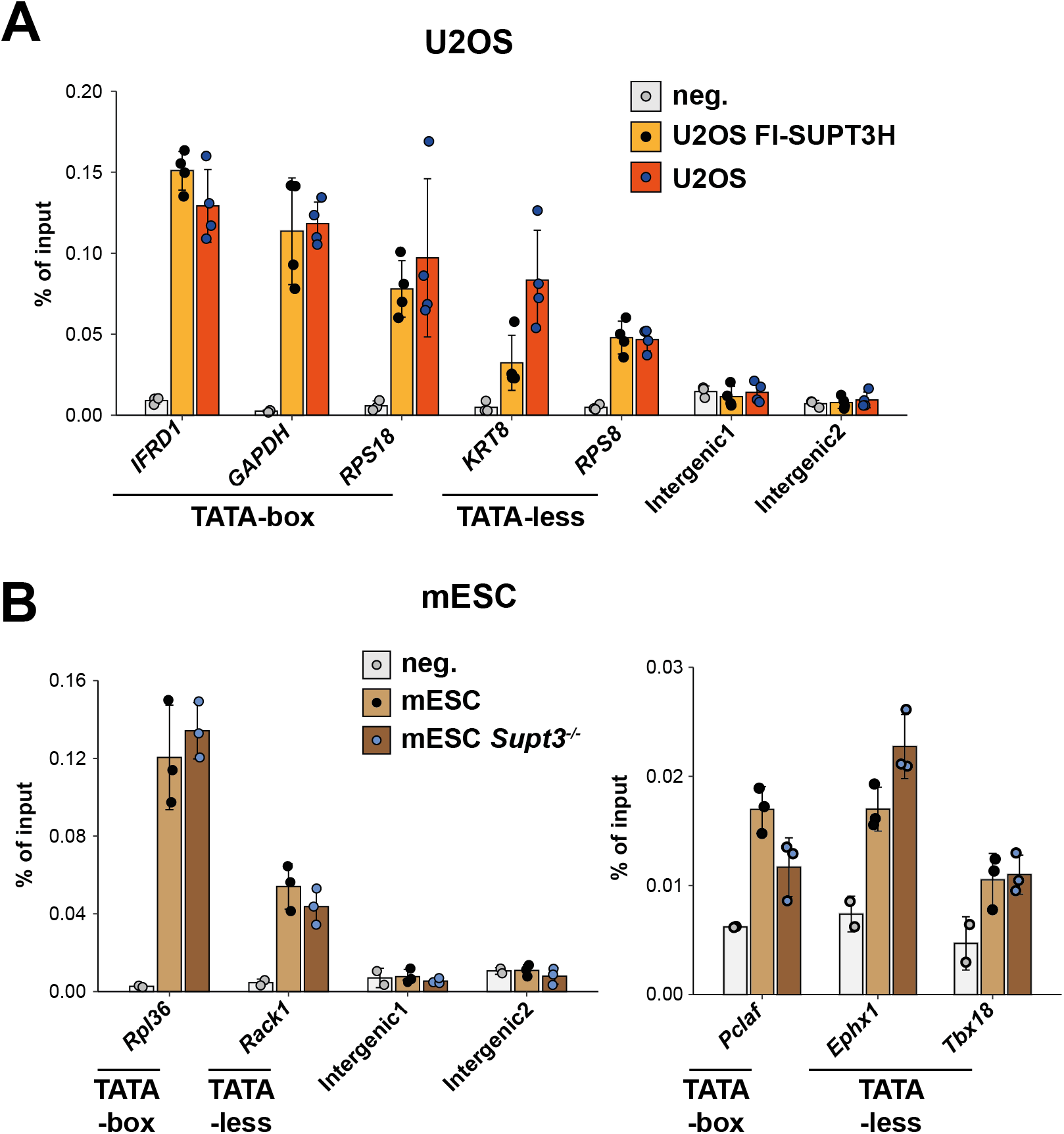
Loss of SUPT3H in human U2OS cells or mouse ESCs does not significantly affect TBP binding at promoters. Quantification of TBP ChIP-qPCR experiments in U2OS and U2OS Fl-SUPT3H cells **(A)** or in WT and *Supt3*^*-/-*^ mouse ESCs **(B)** at promoters of selected genes (as indicated) and at two independent intergenic regions. **(A and B)** TATA-less and TATA-box-containing promoters are indicated. ChIP-qPCR experiments without antibody were used as negative control (neg.). Error bars show mean ± standard deviation (SD) of at four and three biological replicates using two independent clones for human and mouse data, respectively, in each sample the mean shows at least four technical RT-qPCR replicates. Statistical test performed is two-sided Wilcoxon rank-sum test. Statistical significance (*p*<0.05) was not reached.

## Discussion

Previous SAGA subunit deletion and purification studies in *S. cerevisiae* demonstrated that Spt3 is not essential for viability and that ySAGA can assemble in the absence of Spt3, and in the absence of the Spt3/Spt8 TBP interacting module (9,24,25). However, constitutive deletion of *SPT3* has a global effect on Pol II transcription, which decreases the expression of the majority of genes (18).

In our present study we show that SUPT3H is not required for human U2OS cell survival, that mammalian SAGA complexes lacking SUPT3H can stably assemble, and that TBP recruitment at several TATA-containing and TATA-less promoters is not influenced by the absence of SUPT3H in mammalian SAGA. Nevertheless, mouse ESCs lacking SUPT3H show impaired self-renewal capacities and growth potential. Our experiments further show that constitutive loss of SUPT3H in both human U2OS and mouse ESCs grown in FCS+LIF+2i medium affects only a small subset of genes.

Some genes transcriptionally deregulated may play important roles in the cellular homestasis of either U2OS or ESCs lacking SUPT3H: i.e. several histone genes of the *HIST1* cluster, *PAX7* and *JAK3* in U2OS cells), or *Sall1* and *Pclaf* in mESCs (Supplementary Table 6). Further studies will be necessary to identify which effects are directly caused by the loss of SUPT3H and how these changes contribute for example to the growth and self-renewal defects observed in mESCs.

Importantly, *Supt3* deletion in mice indicated that SUPT3H is important for mouse embryogenesis, but not for early development, as embryos die between E9.5 and E14.5 (48). This observation suggests that SUPT3H is not essential for early mouse development, similarly to other SAGA subunits, such as GCN5, PCAF, SUPT20H, USP22 and ATXN7L3 (49-54).

Cryo-EM structural studies suggested that yeast Spt3, or human SUPT3H, maybe an important structural subunit of SAGA, as both double HF-containing proteins assemble with three pairs of other HFD-containing SAGA subunits to form a deformed histone octamer-like structure (10,11,22). Importantly, it has been described in ySAGA that while the six histone-fold pairs in the histone-fold octamer-like structure are oriented similarly as in the canonical nucleosome, the intramolecular HF pair of Spt3 is tilted by 20 degrees compared with its analogous nucleosome histone H2A–H2B histone pair (10). This tilt therefore could almost completely free Spt3 from its association with the histone-fold octamer. However, ySAGA subunit composition is not affected by the loss of Spt3 (9,24,25), which we confirmed in both human and mouse cells. Interestingly, plants are probably also lacking a Spt3 orthologue (5,55). Thus, we speculate that the flexibility of Spt3 may be important for TBP binding and release and perhaps to couple its delivery with other SAGA regulatory activities through an allosteric mechanism.

Moreover, no ortholog of ySpt8 was detected in mammalian SAGA (5), suggesting that the TBP-interacting module described in yeast SAGA, would be affected in mammalian cells lacking SUPT3H. We were therefore surprised that the absence of SUPT3H has a minimal impact on Pol II transcription and TBP recruitment. It is thus conceivable that in mammalian cells the TBP promoter delivery mechanism is less, or not at all, dependent on SUPT3H. Alternatively, other SAGA subunits or TFIID may be able to compensate the constitutive loss of SUPT3H, masking the true *in vivo* contribution of SAGA to TBP delivery. Finally, it is conceivable that SUPT3H may be important for TBP delivery and transcription of genes that are induced in response to specific developmental and/or environmental signals.

## Authors’ contribution

V.F. and E.S. performed most experiments, E.L. generated U2OS-Fl-SUPT3H cell lines, B.M. performed mass spectrometry analysis, D.P. performed the bioinformatics analyses of 4sU-seq experiments. D.H. provided conceptual help, V.F. and L.T. wrote manuscript with support from E.S., B.M., D.P., S.D.V., D.H., and D.D. L.T. conceived and supervised the study.

## Acknowledgements

We thank all members of the Tora lab for protocols, thoughtful discussions and suggestions, A. Ben Shem for comments and critically reading the manuscript, the GenomEast platform [France Génomique consortium (ANR-10-INBS-0009)] for library preparation and sequencing; C. Ebel and M. Philipps for help with FACS sorting, B. Reina San Martin and N. Jung for help with the CRIPR/Cas9 *Supt3* KO experiment, designing and establishing the corresponding vectors, and the IGBMC cell culture facility for help with U2OS, HeLa and E14 ES cells.

## References

1. Roeder, R.G. (2019) 50+ years of eukaryotic transcription: an expanding universe of factors and mechanisms. Nat Struct Mol Biol, 26, 783–791.

2. Chen, X., Qi, Y., Wu, Z., Wang, X., Li, J., Zhao, D., Hou, H., Li, Y., Yu, Z., Liu, W. et al. (2021) Structural insights into preinitiation complex assembly on core promoters. Science.

3. Spedale, G., Timmers, H.T. and Pijnappel, W.W. (2012) ATAC-king the complexity of SAGA during evolution. Gene Dev, 26, 527–541.

4. Helmlinger, D. and Tora, L. (2017) Sharing the SAGA. Trends Biochem Sci, 42, 850–861.

5. Helmlinger, D., Papai, G., Devys, D. and Tora, L. (2021) What do the structures of GCN5-containing complexes teach us about their function? Biochim Biophys Acta Gene Regul Mech, 1864, 194614.

6. Eisenmann, D.M., Arndt, K.M., Ricupero, S.L., Rooney, J.W. and Winston, F. (1992) SPT3 interacts with TFIID to allow normal transcription in Saccharomyces cerevisiae. Genes Dev., 6, 1319–1331.

7. Laprade, L., Rose, D. and Winston, F. (2007) Characterization of new Spt3 and TATA-binding protein mutants of Saccharomyces cerevisiae: Spt3 TBP allele-specific interactions and bypass of Spt8. Genetics, 177, 2007–2017.

8. Grant, P.A., Winston, F. and Berger, S.L. (2021) The biochemical and genetic discovery of the SAGA complex. Biochim Biophys Acta Gene Regul Mech, 1864, 194669.

9. Sermwittayawong, D. and Tan, S. (2006) SAGA binds TBP via its Spt8 subunit in competition with DNA: implications for TBP recruitment. Embo J, 25, 3791–3800.

10. Papai, G., Frechard, A., Kolesnikova, O., Crucifix, C., Schultz, P. and Ben-Shem, A. (2020) Structure of SAGA and mechanism of TBP deposition on gene promoters. Nature, 577, 711–716.

11. Wang, H., Dienemann, C., Stutzer, A., Urlaub, H., Cheung, A.C.M. and Cramer, P. (2020) Structure of the transcription coactivator SAGA. Nature, 577, 717–720.

12. Birck, C., Poch, O., Romier, C., Ruff, M., Mengus, G., Lavigne, A.C., Davidson, I. and Moras, D. (1998) Human TAF(II)28 and TAF(II)18 interact through a histone fold encoded by atypical evolutionary conserved motifs also found in the SPT3 family. Cell, 94, 239–249.

13. Anandapadamanaban, M., Andresen, C., Helander, S., Ohyama, Y., Siponen, M.I., Lundstrom, P., Kokubo, T., Ikura, M., Moche, M. and Sunnerhagen, M. (2013) High-resolution structure of TBP with TAF1 reveals anchoring patterns in transcriptional regulation. Nat Struct Mol Biol, 20, 1008–1014.

14. Han, Y., Luo, J., Ranish, J. and Hahn, S. (2014) Architecture of the Saccharomyces cerevisiae SAGA transcription coactivator complex. EMBO J, 33, 2534–2546.

15. Patel, A.B., Louder, R.K., Greber, B.J., Grunberg, S., Luo, J., Fang, J., Liu, Y., Ranish, J., Hahn, S. and Nogales, E. (2018) Structure of human TFIID and mechanism of TBP loading onto promoter DNA. Science, 362.

16. Gupta, K., Watson, A.A., Baptista, T., Scheer, E., Chambers, A.L., Koehler, C., Zou, J., Obong-Ebong, I., Kandiah, E., Temblador, A. et al. (2017) Architecture of TAF11/TAF13/TBP complex suggests novel regulation properties of general transcription factor TFIID. eLife, 6.

17. Huisinga, K.L. and Pugh, B.F. (2004) A genome-wide housekeeping role for TFIID and a highly regulated stress-related role for SAGA in Saccharomyces cerevisiae. Mol Cell, 13, 573–585.

18. Baptista, T., Grunberg, S., Minoungou, N., Koster, M.J.E., Timmers, H.T.M., Hahn, S., Devys, D. and Tora, L. (2017) SAGA Is a General Cofactor for RNA Polymerase II Transcription. Mol Cell, 68, 130–143 e135.

19. Warfield, L., Ramachandran, S., Baptista, T., Devys, D., Tora, L. and Hahn, S. (2017) Transcription of Nearly All Yeast RNA Polymerase II-Transcribed Genes Is Dependent on Transcription Factor TFIID. Mol Cell, 68, 118–129 e115.

20. Donczew, R., Warfield, L., Pacheco, D., Erijman, A. and Hahn, S. (2020) Two roles for the yeast transcription coactivator SAGA and a set of genes redundantly regulated by TFIID and SAGA. eLife, 9.

21. Yu, J., Madison, J.M., Mundlos, S., Winston, F. and Olsen, B.R. (1998) Characterization of a human homologue of the Saccharomyces cerevisiae transcription factor spt3 (SUPT3H). Genomics, 53, 90–96.

22. Herbst, D.A. E., M.N. Louder,R.K. Dugast-Darzacq,C. Dailey,G.M. Fang,Q. Darzacq,X. Tjian,R. Nogales,E. (2021), Structure of the human SAGA coactivator complex: The divergent architecture of human SAGA allows modular coordination of transcription activation and co-transcriptional splicing, bioRxiv. https://doi.org/10.1101/2021.02.08.430339

23. Stegeman, R., Spreacker, P.J., Swanson, S.K., Stephenson, R., Florens, L., Washburn, M.P. and Weake, V.M. (2016) The Spliceosomal Protein SF3B5 is a Novel Component of Drosophila SAGA that Functions in Gene Expression Independent of Splicing. J Mol Biol, 428, 3632–3649.

24. Wu, P.Y. and Winston, F. (2002) Analysis of Spt7 function in the Saccharomyces cerevisiae SAGA coactivator complex. Mol Cell Biol, 22, 5367–5379.

25. Lee, K.K., Sardiu, M.E., Swanson, S.K., Gilmore, J.M., Torok, M., Grant, P.A., Florens, L., Workman, J.L. and Washburn, M.P. (2011) Combinatorial depletion analysis to assemble the network architecture of the SAGA and ADA chromatin remodeling complexes. Mol Syst Biol, 7, 503.

26. Umlauf, D., Bonnet, J., Waharte, F., Fournier, M., Stierle, M., Fischer, B., Brino, L., Devys, D. and Tora, L. (2013) The human TREX-2 complex is stably associated with the nuclear pore basket. Journal of cell science, 126, 2656–2667.

27. Green, S., Issemann, I. and Sheer, E. (1988) A versatile in vivo and in vitro eukaryotic expression vector for protein engineering. Nucleic Acids Res., 16, 369.

28. Rabani, M., Levin, J.Z., Fan, L., Adiconis, X., Raychowdhury, R., Garber, M., Gnirke, A., Nusbaum, C., Hacohen, N., Friedman, N. et al. (2011) Metabolic labeling of RNA uncovers principles of RNA production and degradation dynamics in mammalian cells. Nat Biotechnol, 29, 436–442.

29. Radle, B., Rutkowski, A.J., Ruzsics, Z., Friedel, C.C., Koszinowski, U.H. and Dolken, L. (2013) Metabolic labeling of newly transcribed RNA for high resolution gene expression profiling of RNA synthesis, processing and decay in cell culture. J Vis Exp.

30. Schwalb, B., Michel, M., Zacher, B., Fruhauf, K., Demel, C., Tresch, A., Gagneur, J. and Cramer, P. (2016) TT-seq maps the human transient transcriptome. Science, 352, 1225–1228.

31. Pfaffl, M.W. (2001) A new mathematical model for relative quantification in real-time RT-PCR. Nucleic Acids Res, 29, e45.

32. Martin, M. (2011) Cutadapt removes adapter sequences from high-throughput sequencing reads. EMBnet.journal, 17, 10–12.

33. Dobin, A., Davis, C.A., Schlesinger, F., Drenkow, J., Zaleski, C., Jha, S., Batut, P., Chaisson, M. and Gingeras, T.R. (2013) STAR: ultrafast universal RNA-seq aligner. Bioinformatics, 29, 15–21.

34. Anders, S., Pyl, P.T. and Huber, W. (2015) HTSeq--a Python framework to work with high-throughput sequencing data. Bioinformatics, 31, 166–169.

35. Love, M.I., Huber, W. and Anders, S. (2014) Moderated estimation of fold change and dispersion for RNA-seq data with DESeq2. Genome Biol, 15, 550.

36. Anders, S. and Huber, W. (2010) Differential expression analysis for sequence count data. Genome Biol, 11, R106.

37. Benjamini, Y., H ochberg, Y. (1995) Controlling the False Discovery Rate: a Practical and Powerful Approach to Multiple Testing. Journal of the Royal Statist. Soc. B 57, 289–300.

38. Dreos, R., Ambrosini, G., Perier, R.C. and Bucher, P. (2015) The Eukaryotic Promoter Database: expansion of EPDnew and new promoter analysis tools. Nucleic Acids Res, 43, D92–96.

39. Muratoglu, S., Georgieva, S., Papai, G., Scheer, E., Enunlu, I., Komonyi, O., Cserpan, I., Lebedeva, L., Nabirochkina, E., Udvardy, A. et al. (2003) Two different Drosophila ADA2 homologues are present in distinct GCN5 histone acetyltransferase-containing complexes. Mol Cell Biol, 23, 306–321.

40. Zybailov, B., Mosley, A.L., Sardiu, M.E., Coleman, M.K., Florens, L. and Washburn, M.P. (2006) Statistical analysis of membrane proteome expression changes in Saccharomyces cerevisiae. Journal of proteome research, 5, 2339–2347.

41. Zhang, Y., Wen, Z., Washburn, M.P. and Florens, L. (2010) Refinements to label free proteome quantitation: how to deal with peptides shared by multiple proteins. Analytical chemistry, 82, 2272–2281.

42. Edgar, R., Domrachev, M. and Lash, A.E. (2002) Gene Expression Omnibus: NCBI gene expression and hybridization array data repository. Nucleic Acids Res, 30, 207–210.

43. Perez-Riverol, Y., Csordas, A., Bai, J., Bernal-Llinares, M., Hewapathirana, S., Kundu, D.J., Inuganti, A., Griss, J., Mayer, G., Eisenacher, M. et al. (2019) The PRIDE database and related tools and resources in 2019: improving support for quantification data. Nucleic Acids Res, 47, D442–D450.

44. Basehoar, A.D., Zanton, S.J. and Pugh, B.F. (2004) Identification and distinct regulation of yeast TATA box-containing genes. Cell, 116, 699–709.

45. de Jonge, W.J., O'Duibhir, E., Lijnzaad, P., van Leenen, D., Groot Koerkamp, M.J., Kemmeren, P. and Holstege, F.C. (2017) Molecular mechanisms that distinguish TFIID housekeeping from regulatable SAGA promoters. EMBO J, 36, 274–290.

46. Eisenmann, D.M., Chapon, C., Roberts, S.M., Dollard, C. and Winston, F. (1994) The Saccharomyces cerevisiae SPT8 gene encodes a very acidic protein that is functionally related to SPT3 and TATA-binding protein. Genetics, 137, 647–657.

47. Belotserkovskaya, R., Sterner, D.E., Deng, M., Sayre, M.H., Lieberman, P.M. and Berger, S.L. (2000) Inhibition of TATA-binding protein function by SAGA subunits Spt3 and Spt8 at Gcn4-activated promoters. Mol Cell Biol, 20, 634–647.

48. Perez-Garcia, V., Fineberg, E., Wilson, R., Murray, A., Mazzeo, C.I., Tudor, C., Sienerth, A., White, J.K., Tuck, E., Ryder, E.J. et al. (2018) Placentation defects are highly prevalent in embryonic lethal mouse mutants. Nature, 555, 463–468.

49. Bu, P., Evrard, Y.A., Lozano, G. and Dent, S.Y. (2007) Loss of Gcn5 acetyltransferase activity leads to neural tube closure defects and exencephaly in mouse embryos. Mol Cell Biol, 27, 3405–3416.

50. Zohn, I.E., Li, Y., Skolnik, E.Y., Anderson, K.V., Han, J. and Niswander, L. (2006) p38 and a p38-interacting protein are critical for downregulation of E-cadherin during mouse gastrulation. Cell, 125, 957–969.

51. Yamauchi, T., Yamauchi, J., Kuwata, T., Tamura, T., Yamashita, T., Bae, N., Westphal, H., Ozato, K. and Nakatani, Y. (2000) Distinct but overlapping roles of histone acetylase PCAF and of the closely related PCAF-B/GCN5 in mouse embryogenesis. Proc Natl Acad Sci U S A, 97, 11303–11306.

52. Atanassov, B.S., Mohan, R.D., Lan, X., Kuang, X., Lu, Y., Lin, K., McIvor, E., Li, W., Zhang, Y., Florens, L. et al. (2016) ATXN7L3 and ENY2 Coordinate Activity of Multiple H2B Deubiquitinases Important for Cellular Proliferation and Tumor Growth. Mol Cell, 62, 558–571.

53. Koutelou, E., Wang, L., Schibler, A.C., Chao, H.P., Kuang, X., Lin, K., Lu, Y., Shen, J., Jeter, C.R., Salinger, A. et al. (2019) USP22 controls multiple signaling pathways that are essential for vasculature formation in the mouse placenta. Development, 146.

54. Wang, F., El-Saafin, F., Ye, T., Stierle, M., Negroni, L., Durik, M., Fischer, V., Devys, D., Vincent, S.D. and Tora, L. (2021) Histone H2Bub1 deubiquitylation is essential for mouse development, but does not regulate global RNA polymerase II transcription. Cell Death Differ.

55. Antonova, S.V., Boeren, J., Timmers, H.T.M. and Snel, B. (2019) Epigenetics and transcription regulation during eukaryotic diversification: the saga of TFIID. Genes Dev, 33, 888–902.

