## Supplemental Figures and Tables for "SUPT3H-less SAGA coactivator can assemble and function without significantly perturbing RNA polymerase II transcription in mammalian cells"

**Running Title:** SUPT3H-lacking SAGA can assemble and function

**Key words:** SAGA (Spt-Ada-Gcn5 acetyltransferase) complex, Spt3, RNA polymerase II, transcription, knock-out, osteosarcoma, mESC, 4sU RNA-seq, histone fold, U2OS cells, mouse embryonic stem cells, proteomic, co-activator, gene regulation.

### Supplemental Figures:

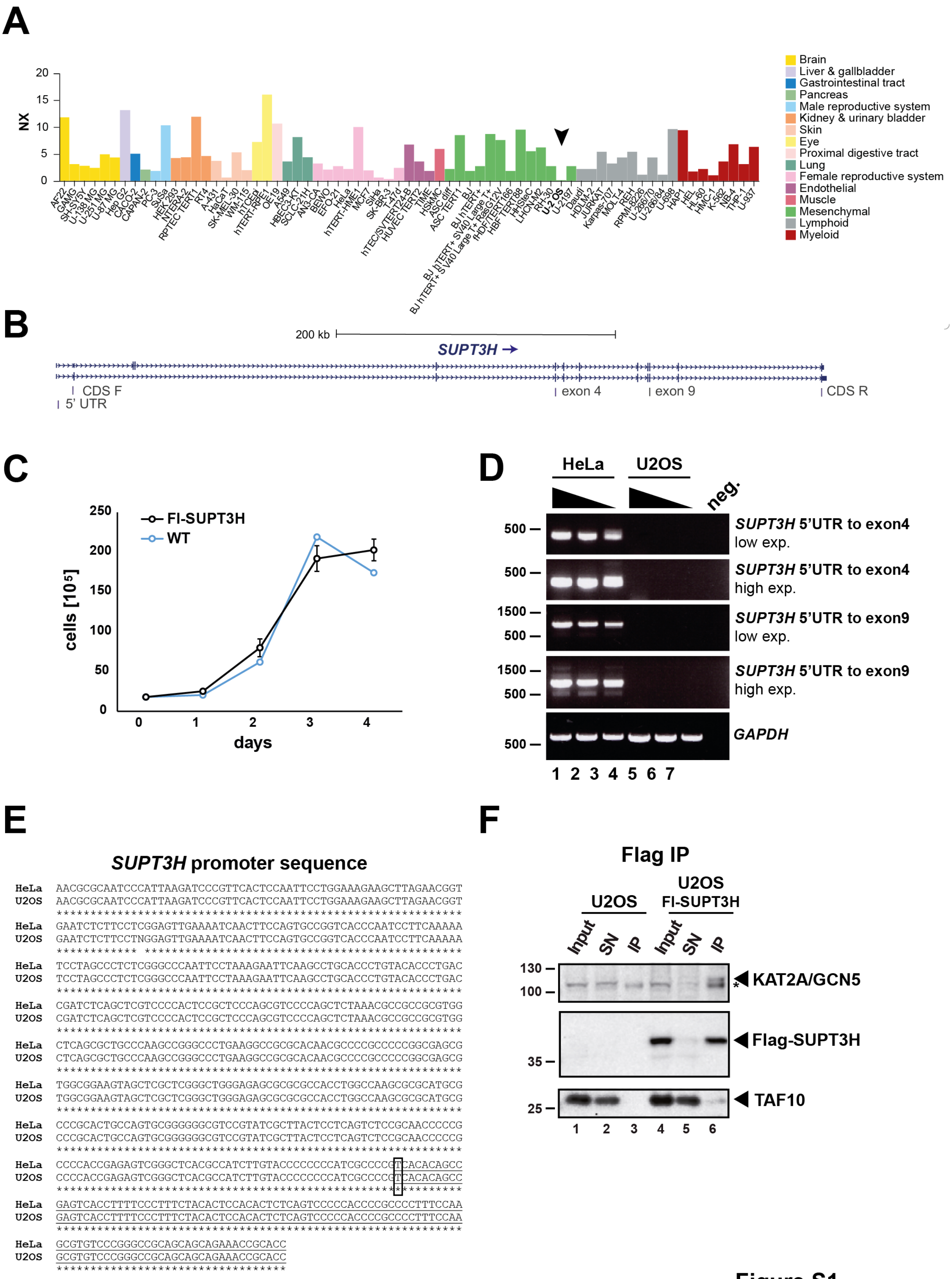


**Figure S1**

**Figure S1: Characterization of *SUPT3H* expression in human U2OS cells. (A)**

Human protein atlas data for *SUPT3H* mRNA levels in cell lines of different origins (as indicated). Arrow indicates human U2OS osteosarcoma cell line. **(B)** Scheme of the two main *SUPT3H* transcript variants showing the position of primers used in Fig. 1A and Supplementary Figure S1D. CDS = coding sequence, R = reverse, F = forward, exp. = exposure. **(C)** Growth curve analysis of WT U2OS cells compared to U2OS-Fl-SUPT3H cells. **(D)** PCR analysis of *SUPT3H* mRNA levels on cDNA obtained from HeLa and U2OS cells. Water served as negative control (neg.) and expression of *GAPDH* was used as a loading control. Low and high exposures (exp.) are shown. **(E)** *SUPT3H* promoter sequence of HeLa and U2OS cells was PCR amplified, sequenced and is represented in 5'-to-3' direction. Underlined sequences represent the 5' portion of the first exon of *SUPT3H*. The transcription start site (TSS) is highlighted by a black rectangle. Stars (*) show perfect homology. **(F)** Western blot assay of input, supernatant (SN) and elution fractions of Flag IPs from U2OS and U2OS Fl-SUPT3H cells using the indicated antibodies.

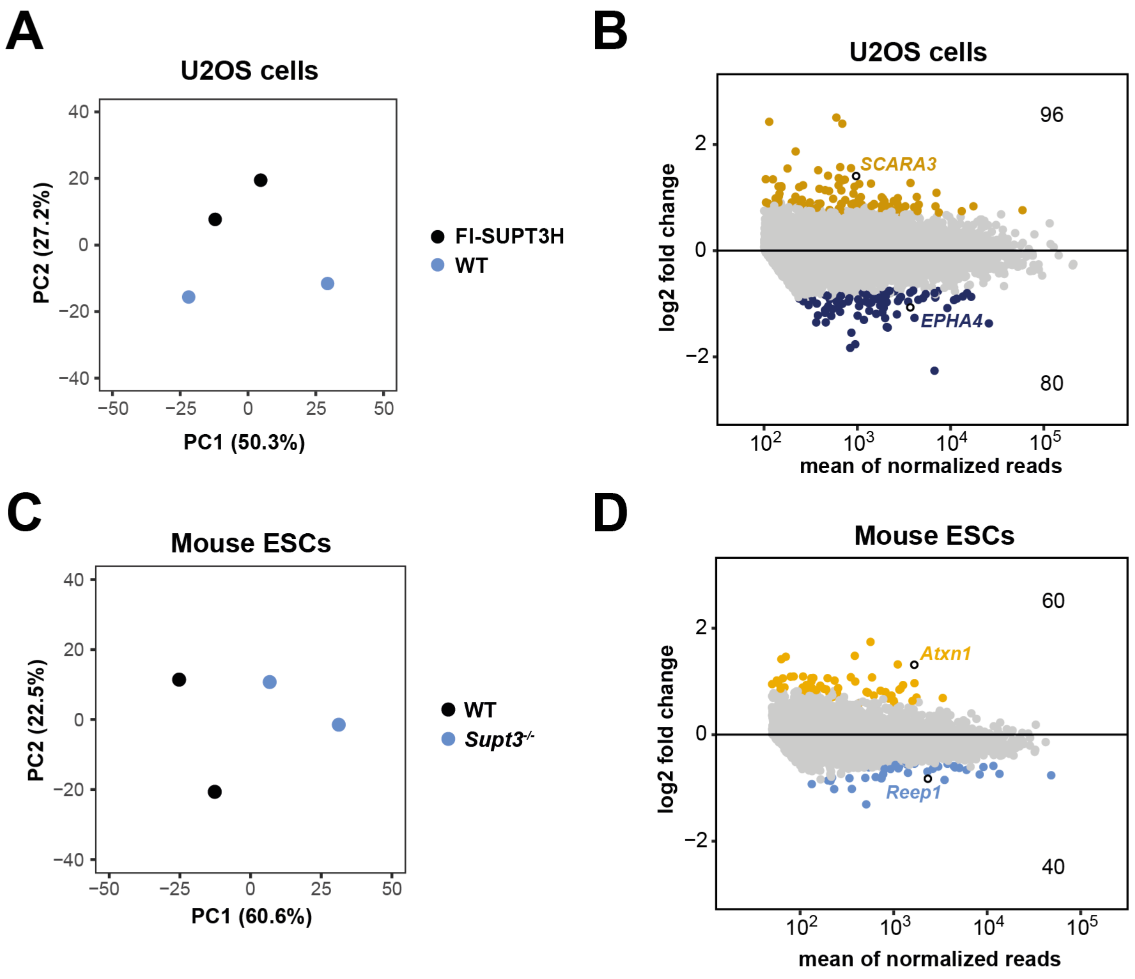


Figure S2

**Figure S2: Additional analysis of 4sU-seq in U2OS and mESC cells. (A)** PCA analysis of 4sU-seq replicates of U2OS WT and U2OS-FI-SUPT3H cells. **(B)** MA plot of 4sU-seq experiments showing log2 fold changes comparing U2OS WT lacking SUPT3H to U2OS FI-SUPT3H cells relative to the mean of normalized reads of U2OS FI-SUPT3H cells with significantly downregulated (log2 fold change < -0.5 and adjusted *p*-value < 0.05) or upregulated (log2 fold change > 0.5 and adjusted *p*-value < 0.05) genes shown in dark blue and dark yellow respectively. The number of significantly upregulated genes is shown in the upper corner, while the number of significantly downregulated genes is shown in the lower corner. A threshold of 100 reads was used to define expressed genes. The position of the two differentially expressed genes shown in (Fig. 2B) are indicated by white circles with black border. **(C)** PCA analysis of 4sU-seq replicates of *Supt3*$^{-/-}$ and WT mESCs. **(D)** MA plot of 4sU-seq experiments showing log2 fold changes between *Supt3*$^{-/-}$ and WT mouse ESCs to the mean of normalized reads of WT mouse ESCs with significantly downregulated (log2 fold change < -0.5 and adjusted *p*-value < 0.05) or upregulated (log2 fold change > 0.5 and adjusted *p*-value < 0.05) genes shown in light blue and yellow respectively. The number of significantly upregulated genes are shown in the upper corner, while the number of significantly downregulated genes is shown in the lower corner. A threshold of 50 reads was used to define expressed genes. The position of the two differentially expressed genes shown in (Fig. 5B) are indicated by white circles with black border.

### Supplementary Table S1: Reagents

| Reagent | Company name | Catalogue # |
|---|---|---|
| Phusion™ High Fidelity polymerase | ThermoFisher Scientific | F-530 |
| NucleoSpin Gel and PCR Clean-up, Mini kit | Machery-Nagel | 740609.50 |
| T4 DNA ligase | Biolabs | M0202 |
| Nucleospin® Plasmid DNA kit | Machery-Nagel | 740588.50 |
| NucleBond® Xtra plasmid purification kit | Machery-Nagel | 740414 |
| Lipofectamine2000 | ThermoFisher Scientific | 11668019 |
| Phire direct PCR kit | ThermoFisher Scientific | F-1265 |
| Foetal calf serum | Sigma Aldrich | F7524 |
| Gentamycin | KALYS | G0124-25 |
| Gelatine solution | Dutcher | P06-20410 |
| Foetal calf serum ES-tested | ThermoFisher Scientific | 10270-106 |
| ʟ-glutamine | ThermoFisher Scientific | 25030-024 |
| β-mercaptoethanol | ThermoFisher Scientific | 31350-010 |
| Penicillin and streptomycin | ThermoFisher Scientific | 15140-122 |
| Non-essential amino acids | ThermoFisher Scientific | 11140-035 |
| CHIR99021 | axon medchem | 1386 |
| PD0325901 | axon medchem | 1408 |
| Paraformaldehyde | Electron Microscopy Sciences | 15710 |
| Alkaline Phosphatase Kit | Vector Laboratories | SK-5100 |
| 4-thiouridine | Glentham Life Sciences | GN6085 |
| TRI® Reagent | Molecular Research Center Inc. | TR 188 |
| 4-thiouracil | Sigma Aldrich | 440736 |
| TURBO DNA-*free*™ Kit | ThermoFisher Scientific | AM1907 |
| RiboPure™ RNA Purification Kit | ThermoFisher Scientific | AM1926 |
| Random hexamer primers | ThermoFisher Scientific | SO142 |
| Transcriptor Reverse Transcriptase | Roche | 03531287001 |
| LightCycler® 480 SYBR® Green 2x PCR Master Mix I | Roche | 04887352001 |
| RNase-free water | Sigma Aldrich | 95284 |
| EZ-link HPDP-biotin | ThermoFisher Scientific | 21341 |
| Dimethy sulfoxid | Sigma Aldrich | D8418 |
| µMACS magnetic beads | Miltenyi Biotec | 130-074-101 |
| RNeasy MinElute Cleanup Kit | Qiagen | 74204 |
| TruSeq Stranded Total RNA LT Sample Prep Kit with Ribo-Zero Gold | Illumina | RS-122-2301 |
| Illumina Stranded Total RNA Prep, Ligation with Ribo-Zero Plus kit | Illumina | 20040525 |
| IDT for Illumina RNA UD Indexes, Ligation | Illumina | 20040553/4 |
| SPRIselect beads | Beckman-Coulter | B23319 |
| ANTI-FLAG® M2 Affinity Gel beads | Sigma Aldrich | A2220 |
| KCl | Sigma Aldrich | P9333 |
| Glycerol | PanReacAppliChem | A0970,5000 |
| Trizma-base | Sigma Aldrich | T1503 |
| EDTA | Euromedex | EU0007 |
| DTT (Dithiothreitol) | Euromedex | EU0006 |
| $MgCl_2$ | Sigma Aldirch | 63068 |
| NP40 (IGEPAL CA-630) | Sigma Aldrich | I-3021 |
| Complete Protease Inhibitor Cocktail, EDTA free | Roche | 11873580001 |
| HCl | VWR | 20252.290 |
| Trichloroacetic acid | Sigma Aldrich | T0699 |
| Urea | Sigma Aldrich | U0631 |
| Iodoacetamide | Sigma Aldrich | I1149 |
| Endoproteinase Lys-C | Wako | 125-05061 |

| | | |
|---|---|---|
| Trypsine | Promega | V5111 |
| Acetonitrile MS grade | Sigma Aldrich | 1207802 |
| Formic acid | Sigma Aldrich | 94318 |

### Supplementary Table S2: Biological Resources.

| **Cell line** | **Description** |
|---|---|
| Human U2OS osteosarcoma cells | ATCC, HTB-96 |
| Human U2OS cells overexpressing Fl-SUPT3H | This study |
| Mouse ES E14tg2a.4 cells | BayGenomics |
| Mouse ES E14tg2a.4 with inactivation of the *Supt3* gene | This study |

### Supplementary Table S3: List of PCR and qPCR primers.

| **Specie** | **Gene** | **Forward** | **Reverse** |
|---|---|---|---|
| Mouse | *mSupt3* genotyping targeted exon | GTGGAAGATGTGGTTCATACACAA | GTGATAAATAGGAGAGCCTGGAAA |
| Mouse | *mSupt3* genotyping spanning deleted region (also used for sequencing) | GTGATCACAGTTCTTTCTTGGAGA | GTGATAAATAGGAGAGCCTGGAAA |
| Mouse | *mSupt3* qPCR targeted exon | GCCCGATGTCTACTACCACT | AGGCCTCCTAGCATCACCTA |
| Mouse | *mSupt3* qPCR untargeted exon | AAAGGCATTGACGAGGATGAC | GATCAAGTCCTGAGCGATCTTC |
| Mouse | *Pou5f1* (Oct4) | CTAGCATTGAGAACCGTGTGAG | GATTGGCGATGTGAGTGATCT |
| Mouse | *Sox2* | GCGGAGTGGAAACTTTTGT | CGGGAAGCGTGTACTTATCCTT |
| Mouse | *Nanog* | CTCCAGCAGATGCAAGAACTC | CTTGCACTTCATCCTTTGGTTT |
| Mouse | *Esrrb* | GAGGACTCCGCCATCAAAT | TAGTGGTAGCCAGAGGCAATGT |
| Mouse | *Tfcp2l1* | ACTACAACCAGCACAACTCTGG | CCCATTCTCAGGAGATAGCTG |
| Mouse | *Klf4* | GTGGGTTAGCGAGTTGGAAA | GTGCAGCTTGCAGCAGTAAC |
| Mouse | *Rn7sk* | TTCCCCGAATAGAGGAGGAC | TGGACCTTGAGAGCTTGTTTG |
| Mouse | *Rpph1* | GGGGGAGAGTAGTCTGAATTGG | CGGAGCTTGGAACAGACTCA |
| Mouse | Intergenic-1 | GAGTGCATGCCATTCCATACAC | GAAACGTCTGCTGTGTGGTAAC |
| Mouse | Intergenic-2 | GTTACTCCATCTTTGCCCCCTAA | ATATGCTTGAAGGACCAGGACAG |
| Mouse | *Rpl36* | CTGTCTCTCTAGCCGGAAAGG | CCCACTTACCTGATGCTCTCC |
| Mouse | *Rack1* | CAAGTATCGTCGTTTCTTCTGC | TTTAATCTCCCGAGAAGCCTTT |
| Mouse | *Pclaf* | GGAGGTACAAATCACGACCACT | TTCACTCTGGACCAATTGACAG |
| Mouse | *Ephx1* | CTGAACAGGGTATTGGCAGTTT | AACGAACTGCTAGGAAGGTGAG |
| Mouse | *Tbx18* | GGTGGGAAGAACTTGCTCATTA | GAGCGGAGTTAGATTCTCCTGA |
| Human | Intergenic-1 | TTGGTGTGAGAGGGTGTAAGTG | GCAGTTAGAGGGCTAAGAATGC |
| Human | Intergenic-2 | TGAAGCAGATCCATTGTCACTT | TTTGGCAGCCTTCCTAATGTAT |
| Human | *GAPDH* | ACGTAGCTCAGGCCTCAAGAC | CTGACTGTCGAACAGGAGGAG |
| Human | *RPS8* | GCAGCTGACACGTAAGTCTCGT | AGATTCACGGTTCGGTTTGTAA |
| Human | *RPS18* | GGTACCCAAGCCATACTCTCATA | ATATAGCTTCCAGGAGGAAGAGG |
| Human | *KRT8* | AAGCTGCTTCTTGGTGAGGAG | CACTCAGGTACGAGGCCTTTC |
| Human | *IFRD1* | GGTTGGGAAGCGCTCAAG | CTAGCGCGGGAGCTAAGG |
| Human | *SUPT3H* CDS | TGAATAATACGGCAGCTAGTCCA | CAGCAGGCTAGAAAAGCCATC |
| Human | *SUPT3H* 5'UTR-exon4 | CGAGTCACCTTTTCCCTTTCTA | TTCAGGAGTGATTACCCTTGCT |
| Human | *SUPT3H* 5'UTR-exon9 | CGAGTCACCTTTTCCCTTTCTA | CTGAATGAAGGTTGCAGAAATG |
| Human | *GAPDH* cDNA | AACGGGAAGCTTGTCATCAAT | GCTGTTGAAGTCAGAGGAGACC |
| Human | *hSUPT3H* PCR primers HA 13662980 and HA13662981 | CCGGCTCGAGATGAATAATACGGCAGC | ATATCCCGGGTCAGCAGGCTAGAAAAGCC |

| | | | |
|---|---|---|---|
| Human | *hSUPT3H* sequencing primers HA 13697387 and HA 13730941 | TGTGCTGTCTCATCATTTTGG | CATGCTCCAGACTGCCTTG |

**Supplementary Table S4: List of sgRNAs used to generate *Supt3*$^{-/-}$ mouse ES E14 cell lines**

| **Name** | **Sequence** | **PAM** |
|---|---|---|
| *mSupt3*-1 | TCCTGAAGCCTGAATTTGGT | AGG |
| *mSupt3*-2 | GTGATGGGATCTATTCAGTG | TGG |

**Supplementary Table S5: List of antibodies used in this study.**

| **Name** | **Company name** | **Catalogue # or reference** |
|---|---|---|
| anti-TBP antibody mouse monoclonal | abcam | ab51841 |
| anti-TAF7 antibody rabbit polyclonal | IGBMC antibody service (3475) | (1) |
| anti-TAF10 antibody mouse monoclonal | IGBMC antibody service (6TA-2B11) | (2) |
| anti-SUPT20H antibody rabbit polyclonal | IGBMC antibody service (3006) | (3) |
| anti-SUPT3H antibody rabbit polyclonal | IGBMC antibody service (3118) | (1) |
| anti-TADA2B antibody rabbit polyclonal | IGBMC antibody service (3122) | This study |
| anti-TADA3 antibody rabbit polyclonal | IGBMC antibody service (2678) | (4) |
| anti-FLAG M2 mouse monoclonal antibody | Sigma Aldrich | F1804 |
| anti-KAT2A/GCN5 antibody mouse monoclonal | IGBMC antibody service (2GC 2C11) | (5) |

**References:**


1. Bardot, P., Vincent, S. D., Fournier, M., Hubaud, A., Joint, M., Tora, L., and Pourquie, O. (2017) The TAF10-containing TFIID and SAGA transcriptional complexes are dispensable for early somitogenesis in the mouse embryo. *Development* **144**, 3808-3818
2. Jacq, X., Brou, C., Lutz, Y., Davidson, I., Chambon, P., and Tora, L. (1994) Human TAFII30 is present in a distinct TFIID complex and is required for transcriptional activation by the estrogen receptor. *Cell* **79**, 107-117
3. Krebs, A. R., Karmodiya, K., Lindahl-Allen, M., Struhl, K., and Tora, L. (2011) SAGA and ATAC histone acetyl transferase complexes regulate distinct sets of genes and ATAC defines a class of p300-independent enhancers. *Molecular cell* **44**, 410-423
4. Orpinell, M., Fournier, M., Riss, A., Nagy, Z., Krebs, A. R., Frontini, M., and Tora, L. (2010) The ATAC acetyl transferase complex controls mitotic progression by targeting non-histone substrates. *The EMBO journal* **29**, 2381-2394
5. Brand, M., Moggs, J. G., Oulad-Abdelghani, M., Lejeune, F., Dilworth, F. J., Stevenin, J., Almouzni, G., and Tora, L. (2001) UV-damaged DNA-binding protein in the TFTC complex links DNA damage recognition to nucleosome acetylation. *Embo J* **20**, 3187-3196.
